# Mature plant chloroplasts form reversible gyroid cubic membranes

**DOI:** 10.64898/2026.06.12.731806

**Authors:** Michał Bykowski, Anna Węgrzyn, Wojciech Wietrzyński, Alicja Bukat, Joanna Wójtowicz, Radosław Mazur, Fanny Le Blanc, Zuzanna Bieńko, Karina Kwapiszewska, Gerd E. Schröder-Turk, Benjamin D. Engel, Łucja Kowalewska

**Affiliations:** Department of Plant Anatomy and Cytology, Faculty of Biology, University of Warsaw, Warsaw, Poland; Biozentrum, University of Basel, Basel, Switzerland; Department of Plant Biochemistry, Faculty of Biology, University of Warsaw, Warsaw, Poland; Institute of Physical Chemistry, Polish Academy of Sciences, Warsaw, Poland; School of Mathematics, Statistics, Chemistry and Physics, Murdoch University, Murdoch, WA, Australia; Department of Materials Physics, Research School of Physics, The Australian National University, Canberra, ACT, Australia

## Abstract

Across kingdoms, cells fold their membranes into precise shapes closely linked to their functions. In mature land-plant chloroplasts, the photosynthetic membranes have been viewed as strictly lamellar and it is unknown whether they can take on a different structure while remaining functional. Here, we show that mature *Arabidopsis thaliana* chloroplasts can transform this network into a gyroid-type cubic membrane, which we call the gyrobody. The gyrobody forms reversibly during the night and preserves photosystem II photochemistry. A decrease in stromal side thylakoid surface charge, caused by lower protein phosphorylation, triggers the lamellar-to-gyroid transition which the curvature-inducing lipid MGDG facilitates. This shows that the mature plant thylakoid network is not locked into its lamellar form, revealing unexpected structural flexibility of this system.

## Introduction

Biological membranes are geometrically diverse. Across kingdoms, cells organize membranes into architectures whose geometry is inseparable from function, and the same fundamental shapes recur in unrelated contexts (*1–3*). Among the most structurally intricate of these are periodic bicontinuous membranes, continuous membrane labyrinths folded into repeating three-dimensional patterns (also referred to as cubic membranes) (*4*). They appear in stressed endoplasmic reticulum, mitochondria and, transiently, during plastid development, yet their functional significance remains largely unresolved (*4–7*). Few systems illustrate the geometry-function coupling more directly than the thylakoid network of plant chloroplasts, a continuous membrane system whose organization underlies solar energy capture and atmospheric oxygen production (*8–10*). In mature plant chloroplasts, thylakoids adopt a single, highly conserved configuration – stacked grana connected by helically wound stroma lamellae in a pitch-balanced arrangement that aligns grana perpendicular to incident light (*11–15*). This lamellar organization has been regarded as the only membrane geometry present in mature chloroplasts of land plants (*8*, *16*).

The lamellar organization of mature plant thylakoids reflects a tight link between structure and photosynthetic function. The lateral segregation of photosystem II (PSII) into stacked grana and photosystem I (PSI) into stroma lamellae enables independent regulation of the two photosystems and balances the distribution of excitation energy between them (*11*, *17–19*). The periodic stacking of these lamellae along a single axis further supports directional light capture through photonic effects (*20*, *21*). Within mature chloroplasts, thylakoids reorganize dynamically in response to changing light conditions and environmental stress, yet every documented rearrangement has preserved the underlying lamellar topology (*8*, *19*, *22*). However, this picture of lamellar uniformity does not apply to plastids in the early stages of plant development. In dark-grown seedlings, etioplasts house a cubic membrane network called the prolamellar body (PLB), which lacks assembled photosynthetic machinery and converts directly into the grana-stroma network upon illumination (*6*, *23–25*). This shows that plant plastid membranes can, in principle, interconvert between bicontinuous and lamellar topologies, though whether a photosynthetically competent membrane can do so remains unanswered. Among other photosynthetic organisms, cyanobacteria and algae adopt more varied thylakoid architectures. These include gyroid-type cubic configurations that form in the green alga *Zygnema* at the end of log-phase growth (*16*, *26*), suggesting that the strict lamellar organization of land plants may be an evolutionary specialization rather than a physical necessity. These considerations prompted us to ask whether fully developed thylakoid networks of land plant chloroplasts can transform into non-lamellar configurations while maintaining photosynthetic function.

Here, we report that mature plant chloroplasts can reorganize their lamellar thylakoid network into a gyroid-type cubic architecture, which we call the *gyrobody*. Gyrobodies form with unit cell (UC) dimensions of ∼500 nm, approximately six times larger than the length scale of the diamond-type PLB. Using a genetic screen along with cryo-electron tomography (cryo-ET) and SPIRE-based structural analysis, we find gyrobodies in *Arabidopsis thaliana* mutants that affect thylakoid protein phosphorylation (*stn7-1*, *stn7-1 stn8-1*) and xanthophyll cycling (*aba1-6)*. We also observe them in wild-type plants under prolonged darkness or in older leaves. By taking advantage of the full reversibility of gyrobody formation during day-night cycles in *stn7-1*, we monitor the lamellar-to-cubic transition through structural, biochemical, and spectroscopic analyses. The results show that a reduced stromal side thylakoid surface charge density, caused by decreased protein phosphorylation, triggers the transformation. Elevated levels of the curvature-inducing lipid monogalactosyldiacylglycerol (MGDG) help facilitate the subsequent gyroid folding. Rather than acting as a developmental intermediate or a pathological response, the gyrobody demonstrates that mature plant thylakoids can take on a cubic minimal surface membrane conformation while retaining full biochemical functionality. This links the geometry of triply periodic surfaces to the structure of photosynthetic membranes and the mobility of soluble compounds in the chloroplast stroma.

## Results

### Mature Arabidopsis chloroplasts contain gyroid-type cubic membranes

To test whether mature chloroplasts can adopt non-lamellar configurations during normal day/night growth, we screened Arabidopsis mutants impaired in thylakoid organization – including protein phosphorylation and dephosphorylation, acetylation, curvature-inducing proteins, light-harvesting antennae, lipid biosynthesis, and carotenoid metabolism (Fig. 1A, fig. S1A–F). Plants were grown under short-day conditions and sampled at the end of the night to ensure that chloroplasts were fully mature and starch-free. Transmission electron microscopy (TEM) was used to reveal bicontinuous membrane arrangements in three backgrounds: *stn7-1*, the *stn7-1 stn8-1* double mutant, and *aba1-6* (Fig. 1A, fig. S2A–B). All three are impaired in thylakoid dynamics. STN7 phosphorylates light-harvesting complex II (LHCII) to enable state transitions, STN8 phosphorylates PSII core proteins, while ABA1 (zeaxanthin epoxidase) participates in both abscisic acid biosynthesis and the xanthophyll cycle.

**Fig. 1.**
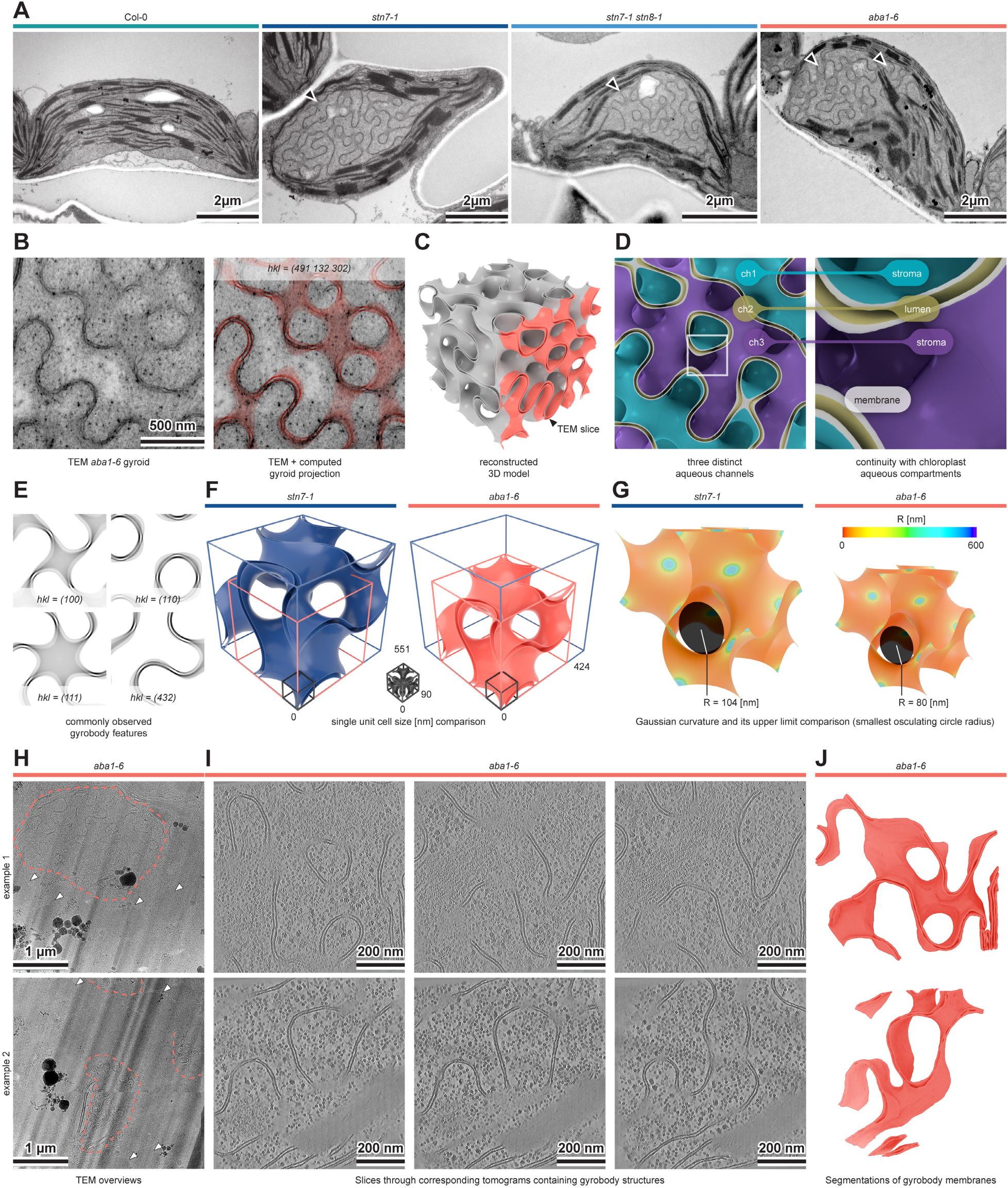
Mature plant chloroplasts can form gyroid-type cubic membranes. (A) Representative TEM images of chloroplasts from Col-0, *stn7-1*, *stn7-1 stn8-1*, and *aba1-6* Arabidopsis plants. Plants were grown for 8 weeks under an 8/16 h light/dark cycle in low light, and samples were collected at the end of the night (16 hD). Cubic membrane structures (black arrowheads) coexist with grana and stroma thylakoids in the mutant chloroplasts. (B) Close-up of an *aba1-6* cubic membrane region (left) and the SPIRE-computed gyroid projection matched to it (right). The computed projection requires slight warping for precise alignment (see Fig. S2). (C) 3D reconstruction of the gyroid structure from SPIRE template matching. The coral-colored region indicates the thickness of the TEM slice used for matching. (D) Close-up of the reconstructed gyroid model with the three aqueous channels color-coded (left), and their connections to the stroma and lumen (right). (E) Computed gyroid projections corresponding to gyrobody features commonly observed in TEM chloroplast cross sections. (F) Reconstructed 3D models of single unit cells (UCs) of gyroids in *stn7-1* (blue) and *aba1-6* (coral) with the largest and smallest observed gyrobody lattice parameter (more examples in Fig. S2). The black UC in the middle shows a PLB unit cell from these mutants for comparison. All UCs are drawn to scale based on SPIRE analysis of TEM data. (G) Gauss curvature distribution of the same UCs as in (F) with osculating circles indicating the most strongly curved points). TEM data from at least three biological replicates per genotype. (H) Two representative TEM overviews of the cryo-lamellae from isolated *aba1-6* chloroplasts. Grana stacks (white arrowheads) are adjacent to unstacked, undulating membranes of the gyrobody (highlighted with a coral line). (I) Two examples (top and bottom rows) of tomograms containing the gyrobody membranes. Three slices spanning the volume of each tomogram are shown to highlight the convoluted membranes (approximately 25 nm spacing in z-plane between slices). (J) Segmentations of the gyrobody membranes from the tomograms shown in (I).

An analysis using nodal surface models (SPIRE) (*23*) identified these structures as double-layer gyroid-type bicontinuous membranes with three distinct aqueous channels (two network-like domains and the space between the two thylakoid membranes, see Fig. 1B–C, fig. S3). We term them *gyrobodies*, extending the PLB nomenclature to a new class of non-lamellar bodies in plant plastids. The gyroid network maintained continuous connections to the surrounding thylakoid system (Fig. 1D, fig. S2C, video S1), and its smallest aqueous channel was connected directly with the thylakoid lumen, preserving the intermembrane distance characteristic of lamellar thylakoids (fig. S2C). On TEM micrographs, the structures are most readily recognized through characteristic projections along specific crystallographic orientations of the gyroid. The appearance of these projections depends on both the structure scale and section thickness (Fig. 1E). UC sizes ranged from 424 to 551 nm across and within genotypes, resulting in corresponding variability in the upper limit of Gaussian curvature (Fig. 1F–G, fig. S3). The gyrobody has an approximately six times larger length scale than the PLB (∼90 nm) and, by virtue of its double layered structure, a different topology than the PLB. Gyrobody formation is hence governed by different molecular mechanisms than those that stabilise the PLB (Fig. 1F–G). Cryo-ET confirmed the gyroid architecture in three dimensions, ruling out a fixation artifact (Fig. 1H-J). The gyrobody regions were seamlessly continuous with flanking domains of typical grana–stroma thylakoid organization, forming a single membrane continuum and indicating partial rather than complete membrane transformation (Fig. 1A, Fig. 1H-J, fig. S2B).

Gyrobodies were restricted to mature chloroplasts of adult, non-flowering plants (fig. S2A–B). Etioplasts and young chloroplasts of *stn7-1* and *aba1-6* contained no gyroid membranes. Only in one-week-old *aba1-6* leaves, chloroplasts showed incipient thylakoid bending (fig. S4A–B). The developmental restriction, together with the difference in length scale and the differing surface type, distinguishes the gyrobody from the PLB both geometrically and developmentally. These results indicate that the thylakoid network of mature plant chloroplasts can adopt a large-length-scale non-lamellar configuration characterized by a gyroid-type surface.

### Gyrobody formation is fully reversible within a single light–dark cycle

Given the extent of membrane reorganization resulting from gyrobody formation, we asked whether the structure represents a terminal state of mature thylakoids or a dynamic, reversible configuration. To address this, we followed gyrobody dynamics across the diurnal cycle in *stn7-1* and *aba1-6*.

In *stn7-1*, gyrobodies formed during the dark period and fully disassembled upon return to light (Fig. 2A–B, fig. S5). In *aba1-6*, by contrast, gyroid structures were constitutively present but varied in size and regularity over the same time course (Fig. 2B, fig. S4C). The full reversibility in *stn7-1* provided a temporal system for tracking the lamellar-to-gyroid transition.

**Fig. 2.**
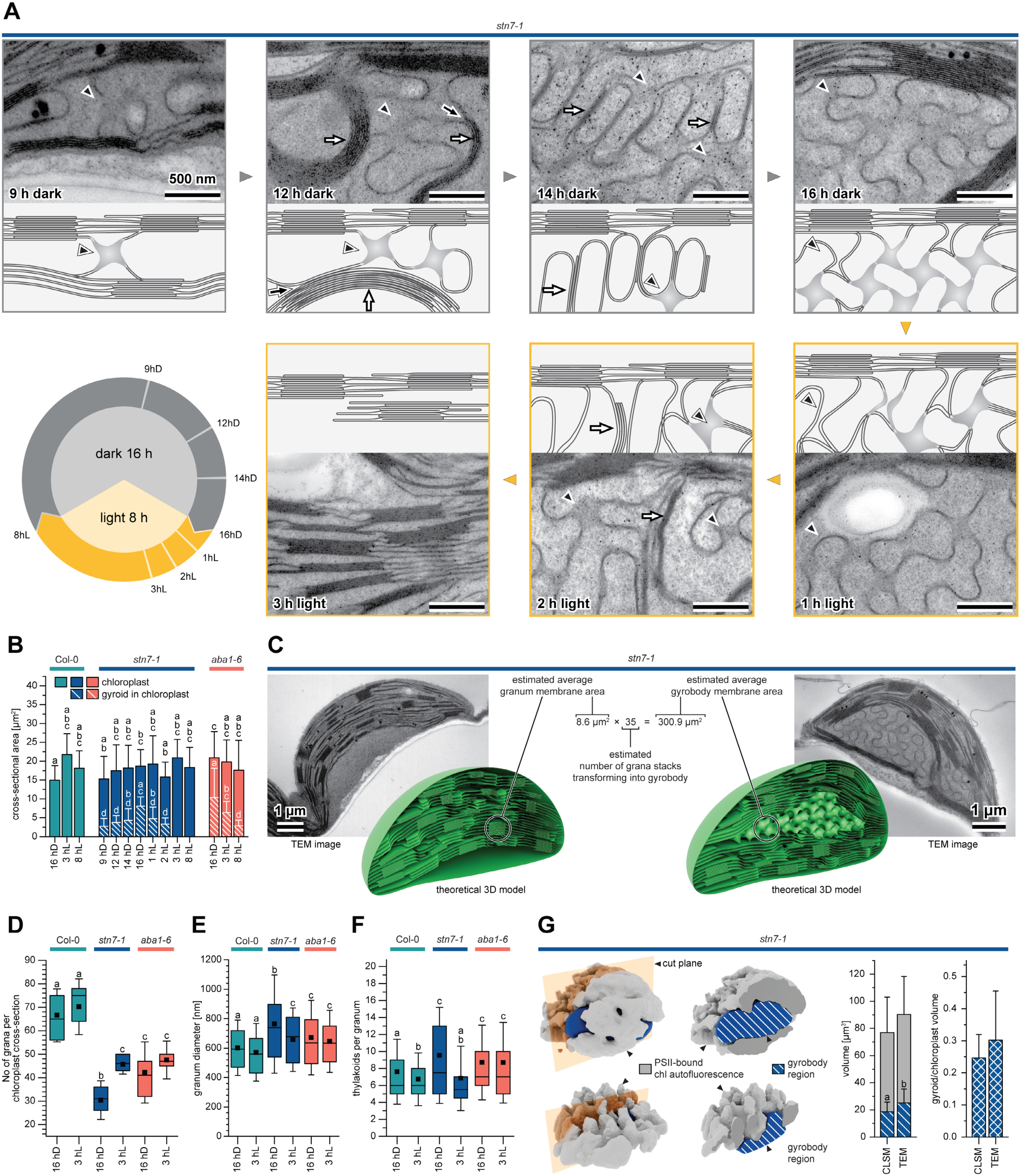
The gyrobody forms and disassembles reversibly during the day/night cycle in *stn7-1* plants through grana stack disintegration. (A) Steps of gyrobody formation and disassembly in *stn7-1* plants at selected time points across the day/night cycle. Each stage is shown as a representative TEM close-up paired with a schematic illustration based on the data. Gyroid features are marked with black arrowheads; grana stacks undergoing transformation are indicated by white arrows. The sampling scheme is shown at the bottom-left. Scale bars: 500 nm. For micrographs of whole chloroplasts, see Fig. S5. (B) Cross-sectional area of chloroplasts (solid bars) and of gyrobodies within chloroplast cross-sections (dashed bars) at selected time points across the day/night cycle in Col-0, *stn7-1*, and *aba1-6* plants. Error bars indicate SD. Different letters indicate significant differences (p < 0.05, Levene’s test for variance homogeneity, subsequently Welch ANOVA with Games-Howell post hoc test). n = [11–81] chloroplasts per condition. (C) Composite schematic showing the estimated number of grana stacks incorporated into the gyrobody in a representative *stn7-1* chloroplast. The schematic combines TEM images and 3D Blender reconstructions. Estimates are based on data in (E)–(G). (D) Box plots of the number of grana stacks visible per chloroplast cross-section. Box edges show the 25th and 75th percentiles; error bars indicate SD; horizontal line and square mark the median and mean, respectively. Statistical analysis as in (B). n = [11-77] cross-sections per condition. (E–F) Box plots of grana stack metrics extracted using GRANA software: (E) average granum diameter, (F) number of thylakoids per granum stack. Box plot description as in (D); statistical analysis as in (B). n = [52-961] grana per condition. (G) Reconstructed chlorophyll autofluorescence surface of *stn7-1* chloroplasts (gray) with the modeled gyroid region (dashed blue) (left). (middle) Bar chart of total chloroplast volume (full bar height) and gyroid volume contribution (blue dashed bar) calculated from CLSM and TEM data. (right) Ratio of gyroid to chloroplast volume from both methods. Error bars indicate SD; statistical analysis (Levene’s test for variance homogeneity, subsequently respectively one-way ANOVA or Welch ANOVA). n = [17-37] chloroplasts per method. TEM and CLSM data are from at least three biological replicates per genotype.

Time-series analysis in *stn7-1* revealed a staged transformation (Fig. 2A, fig. S5). The process initiates with rotation and bending of intergranal stroma thylakoid regions, forming the first gyroid unit. In chloroplast cross-sections that cut grana stacks perpendicular to the thylakoid plane (showing all stacked layers), these initial units appear in the crystallographic (100) projection of the gyroid (fig. S6A). Because TEM sections are thin relative to the gyroid UC, the same unit can appear differently depending on the section’s Z-position within the structure (fig. S6B–D). Grana stacks then rotate and bend, with thylakoid disconnection starting at the edges of each stack. At an intermediate stage, partially unfolded grana stacks align perpendicular to non-transforming stacks, forming a periodic assembly with inter-lamellar distances comparable to the stroma aqueous channel of the final gyroid (Fig. 2A, fig. S5). After 16 hours of darkness (16 hD), the gyrobody reaches its regular structure (Fig. 2A–B, fig. S5). Upon illumination at low light intensity, the gyrobody disassembles within 3 hours (3 hL), restoring lamellar organization through an intermediate stage that again shows perpendicular alignment of restoring grana to non-transforming stacks.

To quantify the scale of this transformation, we estimated the membrane area required to build a typical gyrobody within a single chloroplast. We estimated the gyrobody volume by measuring its cross-sectional area across multiple TEM images and assumed that its shape is an oblate ellipsoid. From the SPIRE modelling, an estimate of the gyroid lattice parameter (*a*) was obtained and consequently the UC volume. Using the known surface area of a gyroid unit cell of lattice parameter *a* and accounting for the two parallel membrane sheets in each unit cell, we estimated the total membrane area contained within the gyrobody (Fig. 2C–F). The calculation indicated that more than 35 typically-sized *stn7-1* grana stacks must disassemble to provide the membrane area observed in the 16 hD gyrobody. Consistent with this estimate, the number of grana stacks per TEM chloroplast cross-section dropped by approximately 30% in the presence of the gyrobody (Fig. 2D). Grana membranes, together with stroma lamellae, therefore serve as the primary structural substrate for gyrobody formation.

We validated the ellipsoid-based gyrobody volume estimates against direct three-dimensional measurements. Using confocal microscopy at room temperature, we imaged chlorophyll autofluorescence in chloroplasts within living mesophyll cells. The gyrobody region showed no detectable fluorescence, similar to stroma thylakoids, and could therefore be identified as the non-fluorescing volume within the scanned chloroplast (Fig. 2G, fig. S7). Absolute volumes differed slightly between the methods, but the gyrobody-to-chloroplast volume ratio was consistent across approaches.

These findings show that gyrobody formation is reversible. The transition proceeds through ordered membrane reorganization in which grana stacks progressively disintegrate and contribute their membrane material to the gyroid assembly.

### Gyrobody formation extends to wild-type plants and preserves PSII photochemistry

Gyrobodies were initially identified in mutants impaired in thylakoid dynamics, where formation depended on chloroplast maturity and required the dark phase of the diurnal cycle. We therefore asked whether the same parameters, independent of mutation, could trigger gyrobody formation in wild-type plants, and whether the gyrobody is compatible with normal photosynthetic function. We further characterized the mutant phenotype by testing how light intensity affects gyrobody size.

Using leaf position within the Arabidopsis rosette as a proxy for tissue age, we examined Col-0 plants after a single 16-hour night. Middle-position leaves (used in all earlier experiments and subsequent biochemical analyses) and young leaves showed no gyroid structures, whereas outer leaves displayed partial membrane reorganization consistent with incipient gyrobody formation (fig. S8A). These outer leaves showed no ultrastructural hallmarks of senescence, ruling out tissue degradation as the cause.

We next tested whether extending darkness beyond the standard photoperiod could trigger gyrobody formation in middle-position Col-0 leaves. After 6 days of continuous darkness, irregular gyroid structures appeared (fig. S8B), confirming that gyrobody formation is not strictly mutant-dependent. In *stn7-1*, gyroid cross-sectional size was maintained between the standard 16-hour night and 3 to 6 days of continuous darkness and grana disassembly remained incomplete (fig. S8B, C).

We also tested whether light intensity during growth affects gyrobody size, using *stn7-1* and *aba1-6,* where gyrobody formation occurs reliably. Fully mature plants grown under higher light intensities formed less developed gyrobodies than those grown under low light (fig. S8B). Plants transferred from high to low light one week before sampling still formed smaller gyroids than those grown continuously under low light (fig. S8B), suggesting that developmental history – not only immediate light conditions – shapes transformation capacity.

To assess whether gyrobody presence affects PSII photochemistry, we compared maximal photochemical efficiency (F_V_/F_M_) between gyroid-containing mutants (*stn7-1* and *aba1-6*) and lamellar-only Col-0 controls at the end of the night period. F_V_/F_M_ values in *stn7-1* and *aba1-6* were comparable to those of wild-type controls (fig. S9A–B). Although a slight but statistically significant decrease in F_V_/F_M_ between wild-type and mutants was observed, the results indicate that the presence of gyrobody does not compromise PSII photochemistry. *aba1-6* plants additionally showed elevated non-photochemical quenching (NPQ). This result is consistent with the zeaxanthin overaccumulation characteristic of this mutant and does not appear as a gyrobody-specific effect (fig. S9C).

Together, these results show that gyrobody formation is accessible to wild-type chloroplasts under prolonged darkness or in older tissues, and that gyroid-containing chloroplasts retain PSII photochemical function comparable to lamellar wild-type controls.

### Gyrobodies remain stable in isolated thylakoid membranes

To identify the molecular factors governing the lamellar-to-cubic transition, we needed to obtain isolated membranes containing gyrobodies for biochemical and spectroscopic analyses. We isolated thylakoid membranes and tested whether the gyrobody persists, using three complementary imaging methods: high-pressure freezing and freeze-substitution (HPF-FS) followed by room-temperature TEM, chemical fixation followed by room-temperature TEM, and cryo-TEM. Imaging by all three methods showed that gyroid structures survived isolation. Some deformation was visible, but the cubic arrangement remained recognizable in buffered solution (Fig. 3A, fig. S9D).

**Fig. 3.**
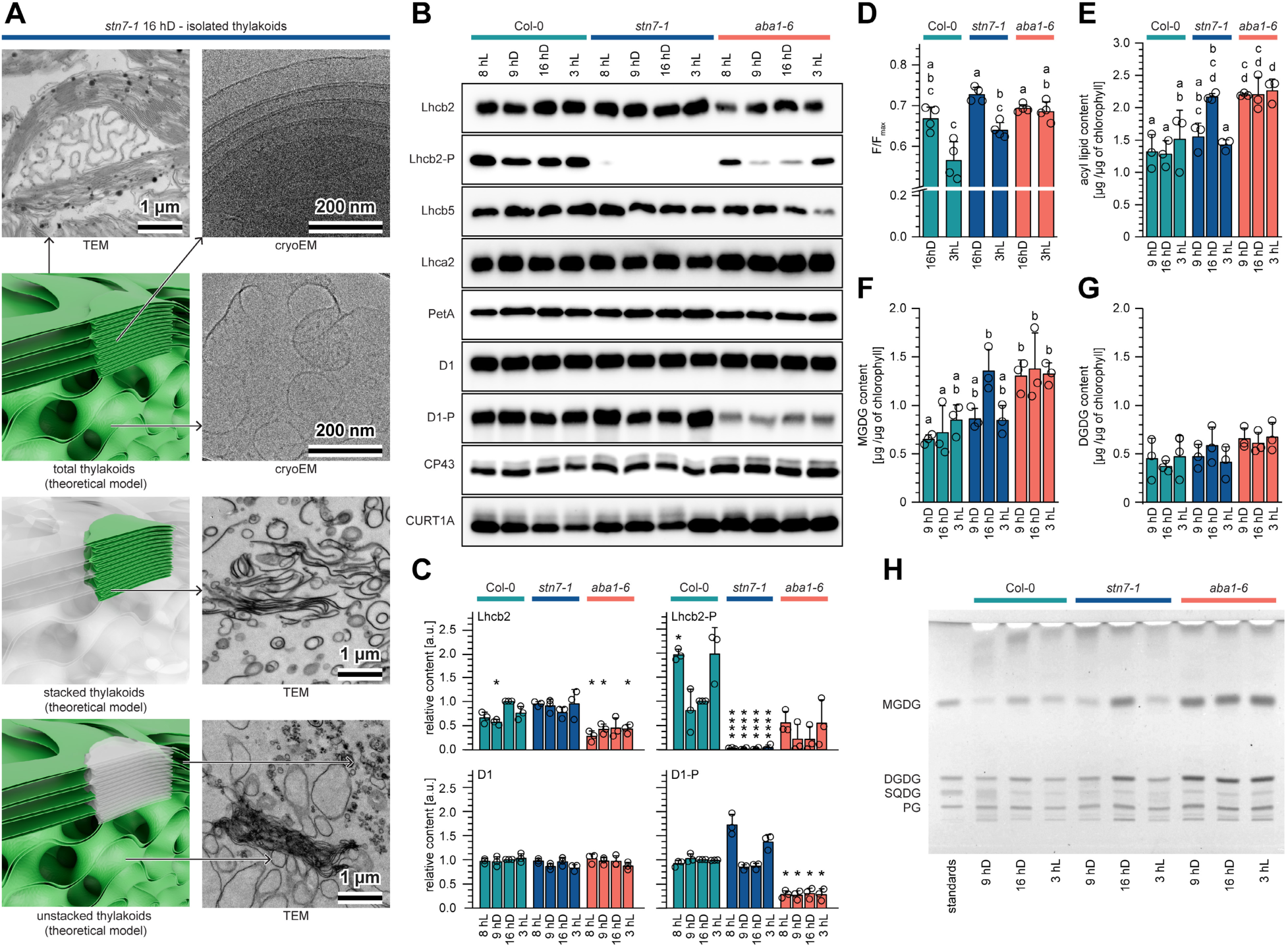
Decreased stromal side thylakoid surface charge initiates gyrobody formation, facilitated by MGDG accumulation. (A) Isolated thylakoids from *stn7-1* 16 hD samples contain fully developed gyrobodies, visualized by TEM (top left) and cryo-TEM (top right and below), with a 3D model indicating the regions shown in the TEM images (black arrows). (bottom) TEM images of thylakoids fractionated by digitonin solubilization, paired with a 3D model showing which fractionated regions (green) correspond to the TEM views (black arrows). For more examples, see fig. S9D-E. (B) Representative immunoblots of selected thylakoid membrane proteins (Lhcb2, Lhcb5, Lhca2, PetA, D1, CP43, CURT1A) and the phosphorylated forms (-P) of D1 and Lhcb2. (C) Densitometric analysis of selected immunoblots from (B); analysis for the remaining proteins in (B) is shown in Fig. S10. Values are normalized to the Col-0 16 hD sample. Error bars indicate SD. Asterisks indicate significant differences from Col-0 16 hD (*: p ≤ 0.05, **: p ≤ 0.01, ***: p ≤ 0.001, ****: p ≤ 0.0001, one-sample t-test). n = 3 biological replicates per antibody. (D) Membrane negative charge estimated by 9-aminoacridine fluorescence; lower F/Fmax values correspond to higher negative charge. Error bars indicate SD. Different letters indicate significant differences (p < 0.05, Levene’s test for variance homogeneity, subsequently Welch ANOVA with Games-Howell post hoc test). n = 6 per condition. (E–G) Acyl lipid content from HPTLC analysis (H), calculated per chlorophyll: (E) total acyl lipids, (F) MGDG, (G) DGDG. Bar description as in (D), statistics (p < 0.05, Levene’s test for variance homogeneity, subsequently, one-way ANOVA with Tukey HSD post hoc test, with no post hoc test needed in (G)). n = 3 biological replicates per condition. (H) Representative HPTLC plate showing the separation of lipid extracts from the analyzed samples; the original plate was in color, shown here in black and white. Densitometric analysis in (E–G) and in Fig. S11 (SQDG, PG) was performed against standard curves prepared for the four analyzed lipid classes; statistical analysis: p < 0.05, Levene’s test for variance homogeneity, subsequently, one-way ANOVA and Tukey HSD post hoc test. Data are from at least three biological replicates.

We also fractionated total thylakoids into stacked and unstacked regions using digitonin solubilization, to see whether the gyrobody would separate into either fraction. The unstacked fraction of *stn7-1* 16 hD samples showed the complex entangled membrane patterns typical of cubic structures (Fig. 3A), while the same fraction from Col-0 showed no such features (fig. S9E). In later experiments, we used either total thylakoids or digitonin fractions, depending on the analysis and on the amount of material needed.

These results show that the gyrobody is stable in isolated membranes and that bicontinuous features are enriched in the unstacked thylakoid fraction.

### Reduced stromal side thylakoid surface charge triggers gyrobody formation, facilitated by elevated MGDG

To identify molecular changes associated with gyroid formation, we exploited the time-dependent changes in *stn7-1*, comparing four time points: end of day (8hL), 9 hours of darkness (9 hD, gyroid initiation), end of night (16 hD, fully formed gyroid), and 3 hours of light (3 hL, lamellar only). *aba1-6* served as a constitutive gyroid-positive control, and Col-0 as a gyroid-negative control.

Western blot analysis showed two phosphorylation patterns, both correlated with gyrobody presence. In *stn7-1*, LHCII phosphorylation was generally absent across all time points, as expected from the loss of STN7 kinase activity. PSII core phosphorylation, however, was reduced specifically at 9 hD and 16 hD – the time points of gyrobody initiation and full formation (Fig. 3B–C). In *aba1-6*, where gyrobodies are constitutive, PSII core phosphorylation was reduced at all time points compared to Col-0. LHCII phosphorylation was also reduced, with the strongest decrease at 9 hD and 16 hD (Fig. 3B–C). ProQ staining confirmed reduced overall thylakoid phosphorylation in *aba1-6* and in *stn7-1* 9 hD and 16 hD samples compared with Col-0 (fig. S10A–C).

Beyond changes in phosphorylation, *aba1-6* also showed reduced LHCII protein levels compared to Col-0 (Fig. 3B-C, fig. S10D). The thylakoid membrane carries a net negative surface charge, largely from the abundant LHCII proteins, which are themselves negatively charged. Protein phosphorylation adds an additional negative charge on top of this baseline. Reduced phosphorylation therefore lowers the membrane’s surface charge density (*27*), and the further reduction of LHCII levels in *aba1-6* would increase this effect. We propose that this neutralization of the negatively charged stromal side of thylakoid membrane triggers lateral splitting of grana stacks and initiates gyrobody formation. Using 9-aminoacridine fluorescence, we measured charge density on both sides of the thylakoid membrane in gyroid-containing samples and in samples with only lamellar thylakoid arrangement (Fig. 3D). This measurement of the total charge did not reveal significant differences between 16hD Col-0 and mutant samples, pointing to the significant role of the luminal side of the membrane in the overall charge values. However, it should be noted that the luminal side of the thylakoids does not play any role in gyroid formation.

Polar lipid analysis normalized to chlorophyll content revealed increased total lipid content in samples with fully developed gyrobodies, mainly due to elevated levels of the curvature-inducing lipid MGDG (Fig. 3E–G, fig. S11A). The MGDG increase was not detectable at the early gyroid formation stage (9 hD in *stn7-1*), suggesting that MGDG accumulation facilitates gyroid folding but is not sufficient to initiate the lamellar-cubic transition. Consistent with this, the *dgd1-2* mutant has an increased MGDG/DGDG ratio but no alterations in protein phosphorylation, and shows characteristic grana bending without their splitting and gyrobody formation (fig. S1).

To test whether other candidates contribute to gyrobody formation, we applied the same reasoning to *aba1-6* that we had used for *stn7-1*. The *aba1-6* mutation affects the xanthophyll cycle and ABA biosynthesis, so we asked whether carotenoid composition or ABA levels change in *stn7-1* gyroid samples. This would mark them as gyrobody-correlated rather than mutant-specific. Carotenoid composition differed in *aba1-6* compared with Col-0 and *stn7-1*, consistent with the zeaxanthin epoxidase deficiency, but no equivalent changes appeared in *stn7-1* gyrobody samples (fig. S11B). Similarly, ABA levels were decreased in *aba1-6* 3 hL plants but not in other gyroid-containing samples (fig. S11C). Membrane fluidity, assessed using Laurdan fluorescence and GP analysis, also showed no differences between genotypes or time points (fig. S11D), indicating that bulk membrane physical properties do not contribute to gyrobody formation.

These results identify reduced thylakoid phosphorylation resulting in the drop in the negative membrane charge at the thylakoid stromal side as the molecular change underlying gyrobody formation. We propose that this neutralization triggers the lamellar-to-cubic transition initiated by grana splitting. MGDG accumulates later and helps fold the released membrane into the gyroid, but does not initiate the transition on its own.

### Gyrobody formation preserves photosystem segregation while loosening protein macro-organization and increasing stromal mobility

We next asked how gyrobody formation affects the arrangement of photosystems in the membrane and the movement of soluble compounds through the surrounding stroma compartment. Low-temperature (77 K) chlorophyll fluorescence of the unstacked thylakoid fraction showed that *stn7-1* at 16 hD has higher PSI and lower PSII–LHCII signals than Col-0 (Fig. 4A–B). *aba1-6*, where gyrobodies are constitutive, showed the same pattern as *stn7-1* at 16 hD, both at the end of the night and after light exposure. This could result from grana unstacking leading to enhanced LHCII-PSI interactions within gyroid structures and increased intensity of the PSI-related band. SDS-PAGE of stacked and unstacked thylakoid fractions of *stn7-1* and Col-0 showed that LHCII proteins are enriched in the unstacked fraction at 16 hD, when gyrobodies are fully formed in *stn7-1* plants (fig. S12). LHCII is typically concentrated in grana stacks, so its increased abundance in the unstacked fraction indicates relocation from the disassembling grana into the gyrobody, while the lateral separation of PSII and PSI is maintained.

**Fig. 4.**
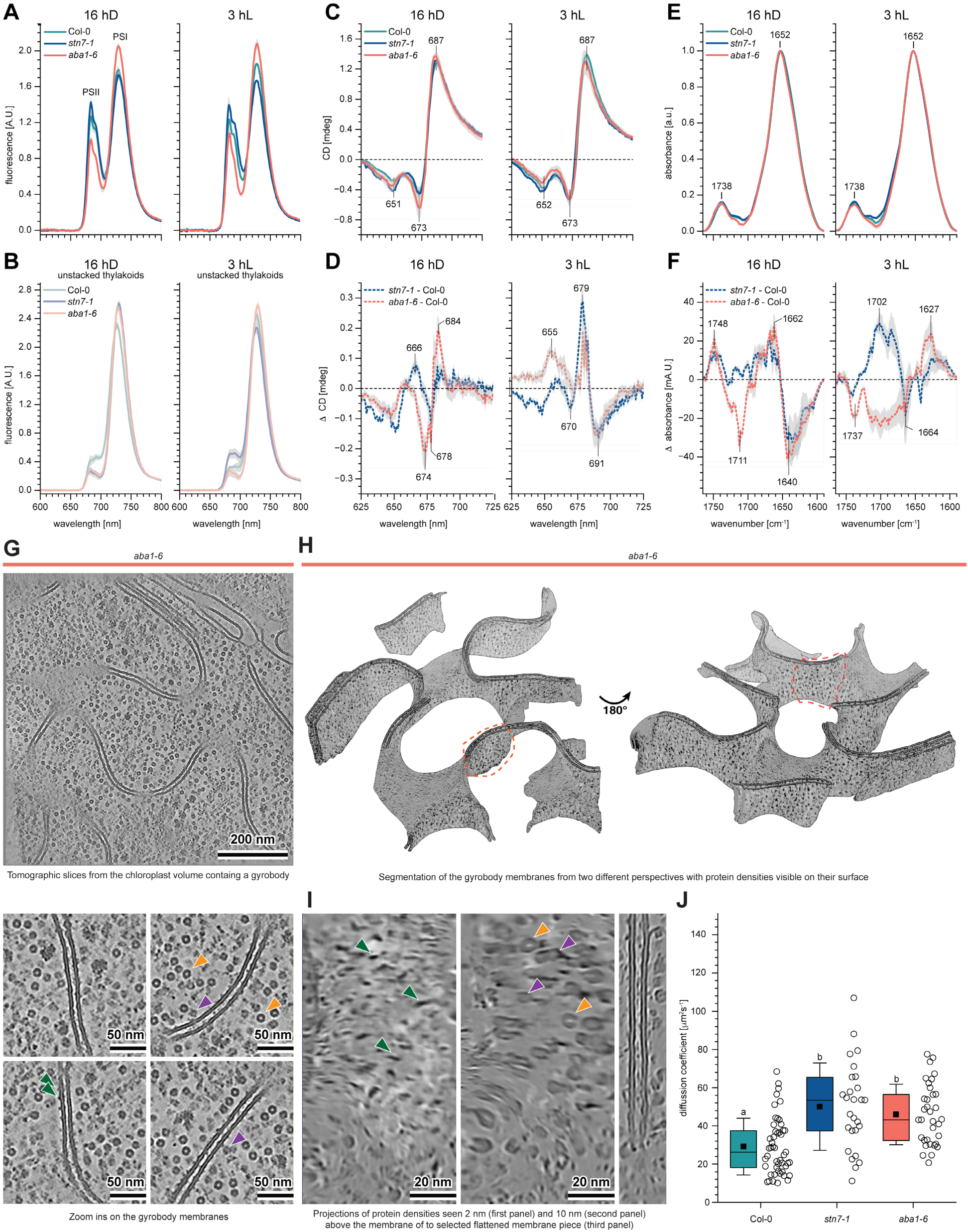
Gyrobody formation preserves photosystem segregation while loosening protein macro-organization and increasing stromal compound mobility. (A–B) Low-temperature (77 K) Chl fluorescence emission spectra (excitation = 440 nm) of total thylakoids (A) and unstacked thylakoids (B), normalized to equal area under the curve. Samples were collected at the end of the night period (16 hD) and after 3 h of light (3 hL). In *stn7-1*, 16 hD samples contain gyrobodies and 3 hL samples have fully lamellar thylakoids, allowing comparison between the two configurations. Col-0 (continuously lamellar) and *aba1-6* (constitutive gyrobody) are shown for comparison. Spectra are averaged across three biological replicates. (C–D) Circular dichroism spectra of total thylakoids (C) and the corresponding difference spectra (D) for 16 hD and 3 hL samples. Spectra were normalized to the maximum absorption of the Chl Q*y* band and averaged across three biological replicates. For full range CD spectra, see Fig. S11E. (E–F) FTIR absorption spectra of 16 hD and 3 hL thylakoid samples (E), normalized to 1 at 1650 cm⁻¹, and the corresponding difference spectra (F), highlighting the amide I and ester C=O regions. Shaded regions in (A–F) represent SD across three biological replicates. (G, top) Tomogram slice showing gyrobody structure. The flattened membranes in the x-y plane are less visible due to lower signal and the missing wedge intrinsic to the cryo-ET acquisition. (G, bottom) Zoom-ins showing four examples of the gyrobody membranes densely decorated with the protein densities. Arrowheads point to the known, photosynthetic complexes: orange - Rubisco, purple - ATP synthase, green - Photosystem I. (H) Membrane segmentation, shown from two perspectives, of the volume shown in (G) with the tomographic densities projected onto its surfaces. Protein densities cover both sides of the structure. Dashed orange outline marks the membrane piece shown in (I). (I) The highlighted membrane piece is shown flattened. Tomographic densities are shown as they are visible 2 nm (left), 10 nm (middle) above the membrane and, as a reference, from the side (right). Photosynthetic complexes are marked with arrowheads as in (G). (J) Diffusion coefficients of the Rhodamine 110 nanotracer in chloroplasts isolated from 16 hD samples containing gyrobodies (*stn7-1* and *aba1-6*) and from chloroplasts with fully lamellar thylakoid networks (Col-0); measurements were targeted to gyrobody or lamellar regions depending on the tested variant. Box edges show the 25th and 75th percentiles; error bars indicate SD; horizontal line and square mark the median and mean, respectively. Different letters indicate significant differences (p < 0.05, Levene’s test for variance homogeneity, subsequently one-way ANOVA with Tukey HSD post hoc test). n = [27-49] chloroplasts per genotype. Data are from at least three biological replicates per condition.

CD spectra showed lower intensity around (−)675 nm in *stn7-1* and *aba1-6* at 16 hD than in Col-0 (Fig. 4C–D, fig. S11E). Bands in this region are associated with chirally ordered LHCII macroassemblies (*28*), and the reduction in their intensities indicates weaker macro-organization of photosynthetic complexes. FTIR of thylakoid samples with fully developed gyrobodies showed reduced absorbance in α-helix bands and in regions linked to membrane-anchored protein interactions (Fig. 4E-F), suggesting looser protein–protein contacts than in lamellar thylakoids.

Cryo-ET of gyrobody-containing chloroplasts revealed the protein distribution within the structure directly. The gyrobody contained no detectable PSII core subunits. Rather, both sides of the gyrobody membranes were densely covered by protein densities, some identifiable as ATP synthases and PSI cores (Fig. 4G-I), which ordinarily are found in the stromal lamellae thylakoids. This suggests that grana disassembly provides the bulk of the gyrobody’s membrane lipid, whereas its protein population resembles that of unstacked thylakoids (PSI, ATP synthase), with PSII retained in the stacks. Because LHCII does not protrude far from the membrane, our cryo-ET analysis cannot directly confirm the LHCII resegregation into the unstacked, gyrobody-enriched regions indicated by the SDS-PAGE and 77 K chlorophyll fluorescence data. It is nonetheless clear that protein complexes densely cover the gyrobody membrane across regions of differing Gaussian curvature, indicating that protein–protein interactions within the curved membrane plane are altered relative to the flat lamellae of typical grana and stromal thylakoids.

Beyond the membrane, gyrobody formation also affected the stromal compartment. Fluorescence lifetime correlation spectroscopy (FLCS) with rhodamine showed that soluble compounds diffuse faster in the gyroid region than in lamellar thylakoid regions of the same chloroplast (Fig. 4J). This likely reflects the more open architecture of the gyrobody: its interconnected channels present fewer obstructions to stromal diffusion than the closely stacked parallel sheets of lamellar thylakoids.

These results show that gyrobody formation reshapes chloroplast molecular organization while preserving photosystem heterogeneity. The lateral separation of PSII and PSI is maintained, but LHCII redistributes out of the grana and the macro-organization of the complexes is weakened. Soluble compounds in the stroma also move more freely in the gyroid region. Whether these changes affect the regulation of photosynthesis under challenging conditions remains an open question.

## Discussion

Our results show that the thylakoid network of mature Arabidopsis chloroplasts can adopt a gyroid membrane configuration. This structure, the gyrobody, forms reversibly during the dark phase of the diurnal cycle in *stn7-1*, is constitutively present in *aba1-6* mutants, and emerges in wild-type plants under prolonged darkness or in older leaves. Until now, thylakoids of mature land-plant chloroplasts were thought to be exclusively lamellar.

Topologically complex, bicontinuous configurations have been visible in plant ultrastructure for decades. Our systematic reanalysis of published electron micrographs (*29*) identifies gyroid-like and other cubic-like membrane arrangements in chloroplasts and chromoplasts across multiple plant families, under conditions ranging from fruit ripening to viral infection to prolonged darkness. In the original publications, these arrangements were not described at all or characterized with terms like curved, sponge-like, or irregular, without being recognized as instances of a common geometric phenomenon. As Linnaeus quoted from Isidore of Seville, *Nomina si nescis, perit et cognitio rerum* — if you know not the names of things, the knowledge of things themselves perishes (*30*). This may explain why gyrobodies in plant chloroplasts were overlooked across decades of chloroplast studies.

### Geometric and topological considerations

The lamellar grana–stroma network and the gyroid architecture are both complex continuous configurations of the thylakoid system, but they differ in how they partition the chloroplast volume. The lamellar architecture creates two aqueous compartments, the lumen and the stroma. The gyrobody creates three channels: the two network-like domains connect to the stroma and the sheet-like domain in between the two membranes connects to the lumen. Whether the transformation between these configurations is a purely geometric reorganization, achievable through continuous deformation, or a topological one requiring local membrane fusion and fission, is not yet clear. Our observation that gyrobody formation begins with thylakoid rotation and grana membrane unstacking does not distinguish between these possibilities. The direct connections between the gyroid small channel and the thylakoid lumen, shown by cryo-ET and TEM, suggest that luminal continuity is preserved throughout the process.

If the transformation does involve a change in topology, intermediate non-minimal configurations are typically required (*31*). Resolving the geometry of the gyrobody transformation will require geometric modeling supported by time-resolved imaging during the transition, for example by cryo-ET or expansion microscopy. This may also clarify the geometric nature of the standard grana–stroma network itself, which has not so far been considered as a possible deformed minimal surface.

### Mechanism of gyrobody formation

We propose a two-stage molecular mechanism for the lamellar-to-gyroid transformation. First, reduced protein phosphorylation lowers the negative surface charge density of the stromal side of the thylakoid membrane, weakening the electrostatic interactions that hold grana stacks together (*27*, *32*). This pattern of reduced phosphorylation is shared between *stn7-1*, which lacks the LHCII kinase STN7, and *aba1-6*, which shows reduced LHCII levels and decreased phosphorylation of both LHCII and PSII core proteins.

Second, elevated levels of MGDG facilitate the negative membrane curvature required for the gyroid surface. MGDG has a small headgroup and a conical molecular geometry. It acts as a curvature-inducing lipid in thylakoid membranes (*33*). The MGDG increase was not detectable at the early gyrobody-formation stage, which means that lipid remodeling helps fold the membrane rather than triggering the transformation. Membrane fluidity did not differ between gyroid-containing and lamellar samples. The transition therefore depends on local molecular features — protein phosphorylation and lipid geometry — rather than on bulk physical properties of the membrane.

*stn7-1* and *aba1-6* differ in which parts of the phosphorylation system remain functional. This explains why gyrobody formation is reversible in the former but constitutive in the latter. *stn7-1* entirely lacks STN7, so LHCII phosphorylation is permanently absent. However, STN8 is still present, and PSII core phosphorylation continues to cycle with the day–night transition. It drops at 9 hD and 16 hD when gyrobodies form and recovers at 3 hL and 8hL. MGDG accumulates later, becoming detectable at 16 hD when the gyrobody is fully formed, and then falls in the light as the structure disassembles. *aba1-6* is different. Both phosphorylation pathways are reduced at all times, LHCII protein levels are lower than in Col-0, and MGDG remains elevated throughout the entire cycle. Without a daily reset, the gyrobody never disassembles.

### Functional implications

The biological function of gyrobodies is not yet clear. The increased mobility of soluble stromal compounds we observed by FLCS suggests one possibility: regulation of metabolite diffusion. The interpenetrating channel systems of the gyroid topology could change how stromal enzymes access thylakoid-associated substrates or products. This may matter most during extended darkness, when chloroplast metabolism shifts from carbon fixation to starch mobilization. Grana disassembly during gyrobody formation also exposes proteins that are normally buried in tightly packed stacks, which could facilitate protein turnover that is otherwise constrained by their inaccessibility to repair and degradation machinery.

Alternatively, gyrobody formation may not itself be adaptive. It may emerge as a default state when the molecular interactions that maintain lamellar organization weaken. Wild-type Arabidopsis plants form gyrobodies under extended darkness, similar to the starvation conditions that trigger cubic membranes in other systems (*34*). In mature chloroplasts, the capacity for the transformation appears to be retained but suppressed under standard growth conditions by active maintenance of membrane surface charge through protein phosphorylation and sufficient amount of LHCII proteins.

Many studies of thylakoid phosphorylation and state transitions rely on biochemical and spectroscopic measurements without accompanying ultrastructural imaging, so effects attributed to molecular changes within a lamellar membrane may, in some cases, reflect a partial shift to a gyroid configuration.

### Evolutionary and technological perspectives

The molecular flexibility we describe in Arabidopsis may extend to land plants more broadly. The wide distribution of these previously unrecognized configurations across plant families (*29*), together with the structural similarity between the gyrobody and the gyroid cubic membranes described in algae such as *Zygnema* sp. (*26*) and *Vaucheria litorea* kleptoplasts (*35*), suggests that the capacity is retained more widely than currently appreciated. Shade-adapted plants and species native to long-night environments are the contexts where this capacity might be most pronounced.

The gyrobody also raises questions in materials biophysics. Its ∼500 nm unit cell is within the visible-light wavelength range, which suggests the possibility of photonic effects in living organisms. In shade-adapted Begonia species, periodically arranged thylakoid lamellae act as a photonic crystal. This setup has been reported to improve light capture at the wavelengths that are most common in shade and increases quantum yield by 5–10% under low light (*36*), providing an example of the use of photonic effects relevant to function of a photosynthetic membrane. In butterflies, convoluted intracellular cubic membranes are observed (*37*) and considered to serve as templates for solid biopolymeric gyroids (*38*, *39*). The resulting single gyroids (*38*, *40*, *41*) have lattice parameter ∼300 nm and produce structural color through photonic interactions. In mammals, Gyroid membranes of similar lattice parameter to the gyrobody have been observed in the mitochondria in the retinal cones of a tree shrew and have been reported to serve an optical function (*42*, *43*). These examples show that gyroid geometries at this scale can interact with light. Because the gyrobody persists during the dark-to-light transition, when light intensity is still low, it may contribute a subtle but significant photonic effect.

With its ∼500 nm unit cell, the gyrobody is substantially larger than other characterized biological cubic membranes, and it has a defined structure and a known biological source. Researchers have created synthetic cubic structures for various technological uses. Lyotropic cubic phases, like cubosomes, are being studied as drug delivery systems (*44*). Block copolymer gyroids are applied in photocatalysts, photovoltaics, and nanoporous membranes (*45*). Most of these synthetic systems have unit cells much smaller than the gyrobody’s ∼500 nm. Bigger unit cells could hold larger cargo than current cubosomes can, such as protein complexes or larger biomolecules. They would also reach the size needed for visible-light photonic effects. Plant-derived components such as MGDG, light-harvesting complexes, and pigments could in principle template biocompatible cubic materials at this scale. The natural gyrobody can switch between lamellar and cubic forms in response to changes in phosphorylation. This suggests that similar reconstituted systems could be made to change states in reaction to external triggers, which would be useful for controlled release.

These applications all depend on a property of the natural structure: the mature thylakoid network is not locked into a single shape but can move between distinct forms. Whether plants make use of this capacity, e.g., for metabolite handling, light capture, or membrane maintenance, and how widely they do so remain open questions. The gyrobody now provides a defined system in which to address them.

## Materials and Methods

### Plant material and growth conditions

The principal *Arabidopsis thaliana* lines analyzed were Col-0 (N1092, *46*)*, stn7-1* (SALK_073254, *47),* and *aba1-6* (N3772, *48*). Additional lines used were indicated in the figures included *tap38-2* (SALK025713), *stn8-1* (SALK_060869), *stn7-1 stn8-1* double mutant (*49*), oe*STN7* (seeds obtained from Pribil Lab; University of Copenhagen), *nsi1/gnat2* (SALK_033944), *dgd1-2* (SAIL_851_G12), *mgd1-2* (SALK_002620), oe*CURT1A* (*50*), *curt1abc* (*50*), *aslhcb2-12* (*51*), *ch1-1* (*52*), *szl1-1 npq1-2* (N66023, *53*), *ccr2-1* (N68150, *54*).

Plants were grown on Jiffy7 peat pellets soaked with mineral medium under short-day conditions: 8 h light at 22°C and 16 h dark at 18°C, with a photosynthetic photon flux density of 30 μmol photons m⁻² s⁻¹ if not stated otherwise. The mineral medium was adjusted to pH 6.0-6.5 and contained 3 mM Ca(NO₃)₂, 1.5 mM KNO₃, 1.2 mM MgSO₄, 1.1 mM KH₂PO₄, 0.1 mM C₁₀H₁₂N₂O₈FeNa, 5 μM CuSO₄, 2 μM MnSO₄·5H₂O, 2 μM ZnSO₄·7H₂O, and 15 nM (NH₄)₆Mo₇O₂₄·4H₂O. Plants were sampled at the end of the light period (8 hL), after 9 h of darkness (9 hD), after 12 h of darkness (12 hD), at the end of the darkness period (16 hD) and after 1 h, 2 h, 3 h and 8 h of reillumination (1 hL, 2 hL, 3 hL and 8 hL respectively). Plant age at sampling was 8 weeks after germination, if not stated otherwise.

Etiolated seedlings were obtained for selected Arabidopsis lines. Seeds were sown in Petri dishes on Murashige and Skoog Basal Medium with Gamborg’s vitamins, solidified with 0.8% Phytagel (Sigma-Aldrich, P8169). Seeds were stratified for 24 h at 4°C and then exposed to 4 h of white light at 120 μmol photons m^−2^ s^−1^ at 23°C to induce germination. Seedlings were subsequently etiolated in darkness at 23°C for 5 days. All samples were harvested in darkness under photomorphogenetically inactive dim green light.

### Chloroplast and thylakoid isolation

All isolation steps were performed at 4°C under dim green light using prechilled buffers. Rosettes were homogenized in buffer A containing 20 mM Tricine-NaOH, pH 7.5, 330 mM sorbitol, 40 mM ascorbic acid, 15 mM NaCl, 4 mM MgCl₂, and 10 mM NaF. Homogenization was performed with a Waring blade homogenizer using 3 short 3 s pulses. The homogenate was filtered through four layers of 50 μm Miracloth and centrifuged at 2000 × g for 4 min at 4°C.

Pelleted chloroplasts were resuspended with a paintbrush in buffer B containing 20 mM Tricine-NaOH, pH 7.5, 15 mM NaCl, 4 mM MgCl₂, and 10 mM NaF to rupture chloroplast envelopes. Thylakoid membranes were collected by centrifugation at 6000 × g for 10 min at 4°C, resuspended in buffer C containing 20 mM HEPES-NaOH, pH 7.0, 330 mM sorbitol, 15 mM NaCl, 4 mM MgCl₂, and 10 mM NaF, centrifuged again at 6000 × g for 10 min at 4°C, and homogenized in buffer C with a Potter-Elvehjem homogenizer (*55*). Buffers were pH-adjusted at 4°C, and NaF was added freshly before use.

### Chlorophyll determination

Chlorophyll concentrations were determined spectrophotometrically after extraction in 80% (v/v) acetone. Absorbance spectra were recorded from 800 to 350 nm, and chlorophyll *a*, chlorophyll *b*, and their sum were calculated from absorbance at 663, and 645 nm using the equations of (*56*).

### Digitonin fractionation of thylakoid membranes

Thylakoid membranes were adjusted to 0.06 mg ml⁻¹ chlorophyll in buffer C and incubated with 0.1% (w/v) digitonin from a freshly prepared 4% (w/v) stock solution, in a final volume of 11 ml in 15 ml Falcon tubes. Samples were rocked at approximately 1 rotation s⁻¹ for 30 min at 4°C in the dark. Samples were centrifuged at 1000 × g for 5 min at 4°C to remove unsolubilized membranes. The supernatant was diluted fivefold with buffer C and centrifuged at 40,000 × g for 30 min at 4°C to collect the stacked-membrane fraction. The remaining supernatant was centrifuged at 100,000 × g for 2 h at 4°C to collect the unstacked-membrane fraction (*57*, *58*). Fraction identity was confirmed by transmission electron microscopy as described below.

### Transmission Electron Microscopy (TEM)

Leaf samples were collected at the indicated time points. The middle vascular bundle was removed, and approximately 2 × 2 mm rectangles were cut from the leaf blade. Samples were fixed at ambient pressure in 2.5% (v/v) glutaraldehyde in 50 mM cacodylate buffer, pH 7.4, for 2 h at 23°C. Samples were washed three times for 10 min in cacodylate buffer and postfixed overnight at 4°C in 2% (w/v) OsO₄ in 50 mM cacodylate buffer, pH 7.4. OsO₄ was prepared from a 4% (w/v) aqueous stock solution. Samples were washed twice for 10 minutes in 50 mM cacodylate buffer, pH 7.4, then twice for 10 minutes in MilliQ water. Sample dehydration was performed in acetone using 10% (v/v) for 10 min, 20% (v/v) for 10 min, 40% (v/v) for 15 min, 60% (v/v) for 30 min, 80% (v/v) for 40 min, and 100% for four 40 min exchanges followed by overnight incubation.

Leaf samples were infiltrated with Agar 100 epoxy resin using 3:1 (v/v) acetone:resin for 4 h, 1:1 (v/v) acetone:resin for 4 h, 1:3 (v/v) acetone:resin for 20 h, pure resin for 24 h, and resin with polymerization accelerator for 2 h. Resin was polymerized for 12 h at 30°C and 48 h at 60°C.

Isolated thylakoid membranes and membrane fractions in quantities equivalent to 100 μg chlorophyll were fixed at ambient pressure in 2.5% (v/v) glutaraldehyde in 50 mM cacodylate buffer, pH 7.4, for 2 h at 23°C. Samples were washed two times for 15 min in cacodylate buffer and postfixed at 4°C in 2% (w/v) OsO₄ in 50 mM cacodylate buffer overnight. Samples were washed twice for 10 minutes in 50 mM cacodylate buffer, pH 7.4, then twice for 10 minutes in MilliQ water. Samples were dehydrated in acetone using 60% (v/v)for 30 min and 100% overnight. Samples were centrifuged between each reagent change 5000 × g.

Fixed membrane isolates were then suspended in small amount of 100% resin with polymerization accelerator and polymerized for 12 h at 30°C and 48 h at 60°C

Ultrathin 70 nm sections were cut with a Diatome diamond knife on a Leica UCT ultramicrotome and mounted on nickel grids. Images were acquired on a JEM 1400 transmission electron microscope (JEOL, Japan) equipped with an 11 Mpix TEM Morada G2 camera (EMSIS GmbH, Germany).

### Thylakoid membrane morphometrics

Grana nanomorphological parameters were measured automatically using the GRANA software (*59*), an AI-based tool that detects grana stacks on calibrated TEM micrographs and returns a set of grana structural parameters. For each experimental variant, TEM images of identical magnification were processed in a single batch in fully automatic mode, followed by hybrid-intelligence filtering: granum masks and orientation assignments were visually verified on the GRANA contact-sheet output, and incorrectly segmented or misoriented stacks were excluded prior to downstream analysis.

Chloroplast and gyrobody cross-sectional areas were measured manually in ImageJ (*60*) by outlining each structure with the polygon-selection tool on calibrated micrographs. The number of grana stacks were counted on cross-sections of entire chloroplasts, and gyrobody cross-sectional dimensions - width and height - were measured directly from TEM micrographs where the entire gyrobody was visible.

### High-Pressure Freezing and Freeze Substitution (HPF-FS)

Leaf samples of approximately 2 × 2 mm were excised from the central leaf blade, infiltrated with 1-hexadecene, mounted in 3 mm aluminum carriers, and frozen at 2100 bar using a Leica HPM100. Freeze substitution was performed in a Leica AFS2 in 1% (w/v) OsO₄ in acetone at −90°C for 20 h. Samples were warmed to −80°C over 4 h, to −60°C over 4 h, held at −60°C for 4 h, warmed to −30°C over 6 h, held at −30°C for 2 h, and warmed to 20°C over 2 h. Samples were washed in 100% acetone and embedded in resin as described above.

### Cryo-Transmission Electron Microscopy with sample preparation

Freshly isolated total thylakoid membranes were washed in buffer D containing 20 mM HEPES-NaOH, pH 7.0, 165 mM trehalose, 7.5 mM NaCl, and 2 mM MgCl₂, and adjusted to 2 mg ml⁻¹ chlorophyll. Lacey carbon or holey carbon-coated R 2/1 copper grids (Quantifoil) were glow discharged for 60 s at 25 mA and 0.38 mbar using a Pelco EasiGlow. Grids with material applied were vitrified by plunge freezing in liquid ethane using a Vitrobot Mark IV operated at 4°C and 100% humidity with 7 s blotting at blot force 7.

Cryo-TEM images were acquired on a Glacios 200 kV cryo-transmission electron microscope equipped with a Falcon 3EC camera operated in linear mode at the Cryomicroscopy and Electron Diffraction Core Facility, Centre of New Technologies, University of Warsaw. Images were recorded at 45,000× with a total dose of 15 e⁻ Å⁻², 3 μm defocus, and 0.31 nm pixel size, or at 73,000× with a total dose of 30 e⁻ Å⁻², 3 μm defocus, and 0.19 nm pixel size.

### Plunge-freezing for Cryo-ET

Vitrification of the samples was done using a Leica GP2 automatic plunger (Leica Microsystems) with the blotting chamber set to room temperature and 100% humidity. 4.2 µL of the sample was placed onto glow-discharged R2/1 carbon-coated 200-mesh copper EM grids (Quantifoil Micro Tools), a small 0.5 µL drop of the buffer was applied on the back side of the grid to improve blotting. Excess sample liquid was then immediately blotted away from the grids for 2-2.5 seconds from the back side. Finally, grids were plunge-frozen in liquid ethane cooled with liquid nitrogen to-182°C. Prepared grids were clipped into “auto-grid” support modified with a cut-out on one side to improve FIB milling access (Thermo Fisher Scientific) and stored in plastic boxes in liquid nitrogen.

### Focused Ion Beam milling

Cryo-FIB milling was performed using an Aquilos dual-beam FIB/SEM instrument (Thermo Fisher Scientific) (*61*). Grids were coated with a layer of organometallic platinum using a gas injection system to protect the sample surface. The samples were milled at shallow angle with a gallium ion beam to produce ∼100-150 nm-thick lamellae. Grids were then transferred in liquid nitrogen to the TEM microscope for tomographic data acquisition.

### Cryo-ET data acquisition

Tilt series (tomographic data) were collected on a 300kV Titan Krios G4 microscope equipped with a Falcon 4i direct electron detector and a SelectrisX energy filter (Thermo Fisher Scientific). Tilt series were acquired using “Tomography” software (Thermo Fisher Scientific) with a dose symmetric scheme starting at the pretilt of-10°, with 2° increment between tilts and spanning a total of 114° (*62*). Data was recorded in EER mode with the pixel size of 1.98 Å/px, constant exposure time, dose per tilt set to between 2.1 and 2.3 e-/Å and target defocus in the range from-2.5 to-4 µm. The total accumulated dose for the tilt-series was kept at approximately 120 e-/Å^2^.

### Tomogram reconstruction and processing

Tilt-series were processed within the SCIPION environment (*63*). Raw frames were aligned using MotionCor2 (v.1.4.0) (*64*). Bad tilts were manually removed. The assembled tilt-series were aligned with patch tracking using the Etomo program from the IMOD (v.4.11.1) software package (*65*). Tomograms were reconstructed by weighted back projection in IMOD (*66*). Tomograms were then denoised with missing wedge inpainting (to improve membrane segmentation) for postprocessing and visualization using ICECREAM or IsoNet2 packages (*67*, *68*).

### Membrane segmentation

Membrane segmentation was performed using MemBrain-seg program from the MemBrain v2 package (*69*) using the publicly available model trained on spinach chloroplast thylakoids. For visualization segmentations were cleaned in Amira3D software (Thermo Fisher Scientific) and rendered and finally images were recorded in UCSF ChimeraX. The flattening of the membrane patch presented in Figure 4 (I) was done using Mpicker software (*70*).

### SPIRE modeling and membrane geometry analysis

The gyroid surface is represented by the nodal approximations (*71*) which are a good approximation of the truly minimal triply periodic gyroid surface (*72*) and which allow for the easy construction of gyroid structures with volume fraction different from 50%.

Triply periodic membrane models based on this representation were generated using SPIRE (*23*), a software that constructs simulated TEM-like projections by representing the membrane as level sets *t* − *t*_2_ < *φ*(*r*) < *t* + *t*_2_ using gyroid approximations by the nodal function 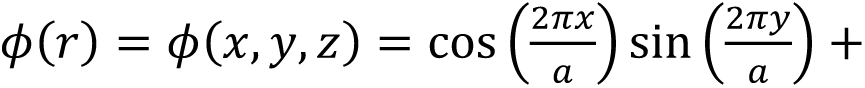 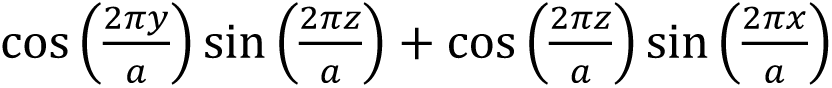 for user-set values of lattice parameter *a*, crystallographic section-plane orientation [ℎ*kl*], and section thickness and membrane thickness (both being functions of the thresholds *t* and *t*_2_)

Model sections were fitted visually to TEM images using reference galleries spanning crystallographic orientations, unit-cell dimensions, and volume proportions. Section thickness was fixed at 70 nm (matching the TEM ultrathin-section thickness) and membrane thickness at 10 nm (matching the apparent osmium-stained thylakoid membrane thickness). Final model slices were overlaid on TEM images in multiply blend mode in Adobe Photoshop and three-dimensional renderings were generated from SPIRE mesh outputs in Blender. Unit-cell dimensions and percolation thresholds (diameters of largest spheres that can traverse the network domains representing the stroma channels) were calculated using SPIRE. For all gyrobodies analyzed here, the best-fit level-set parameter was *t* = 0 (corresponding to balanced double membranes centered on the minimal surface and with equal volume stroma channels).

The Gaussian curvature of the gyroid minimal surface (which sits equidistant between the two bilayers of the gyrobody double membrane) provides a quantification of the membrane curvatures. As a minimal surface (with mean curvature *H* = 0 at every point) this minimal surface is a symmetric saddle with the two principal curvatures at each point equal in absolute value but of opposite sign, *κ*_1_ = −*κ*_2_, so 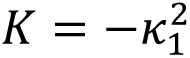 and 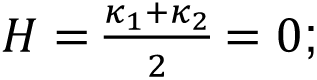; consequently, the radius of curvature is 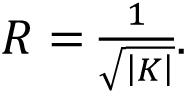. The Gauss curvature is variable across the surface and is strictly negative except at the flat points where *K* = 0. At the most curved point on the surface (indicated in Fig. 1G), the Gauss curvature adopts its largest negative value largest absolute value 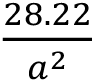 where *a* is the lattice parameter (of the Ia3d space group); this specific value can be determined from the Weierstrass parameterisation of the minimal surface (*73–75*). The surface area of a translational unit cell of the Gyroid minimal surface is *A* ≃ 3.019*a*^2^. A parallel surface offset from the minimal surface by ±*d* has an area *A*(*d*) ≃ 3.019 *a*^2^ + 2*πχd*^2^where *χ* = −8 is the Euler-Poincare index of the Gyroid unit cell (*73–76*). Therefore, the total membrane area in a unit cell of size is *A*_unit_ = 2 × (3.019 *a*^2^ + 2*πχd*^2^) with lattice parameter 424 nm < *a* < 551 nm and the membrane-membrane spacing (the thickness of the lumen compartment) given by 2*d* ≈ 30 nm; The factor 2 reflects that there are two parallel membranes. For *a* = 424 nm, this gives *A*_unit_ = 1.06 μm^2^ and for *a* = 551 nm *A*_unit_ = 1.81 μm^2^. The nodal surfaces described above provide a good but not exact approximation of the Gyroid minimal surface; their Gauss curvature can be calculated using the analytical expression for level set surfaces using the bordered-Hessian determinant formula (*77*); the nodal surface Gauss curvature values in Fig 1G were obtained using that formula and approximate the minimal surface curvatures which were used to calculate *R*.

### Confocal Laser Scanning Microscopy (CLSM)

Samples of single mesophyll cells were prepared as described in (*78*) and imaged using a Nikon A1 MP confocal microscope equipped with a Plan Apo TIRF 100× oil-immersion DIC H objective (NA = 1.45). Chlorophyll autofluorescence was excited at 561.2 nm using a laser operating at 4% of nominal power, and emission was collected from 662 to 737 nm using a 1 Airy-unit pinhole. Z-stacks of 512 × 512-pixel images were acquired with a 60 nm z-step, with detector gain set to 85 and offset to −40 to avoid pixel oversaturation. Collected datasets were deconvolved using AutoQuant X3 software.

### Bright field Light Microscopy (LM)

Semi-thin sections were cut from the same epoxy-resin-embedded blocks used for transmission electron microscopy on a Leica UCT ultramicrotome and mounted on glass slides. Sections were stained with 0.5% (w/v) toluidine blue in water rinsed in distilled water and air-dried. Sections were examined on a Zeiss AxioLab A1 microscope equipped with a Zeiss A-Plan 40× Ph2 objective (NA 0.65).

### Gyrobody volume and membrane-area estimation

Gyrobody dimensions were measured manually from calibrated TEM images in ImageJ Gyrobody volume was approximated as an ellipsoid, 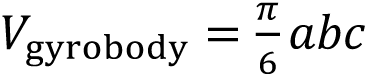 where *a* and *b* are the two ellipsoid axes measured directly from the TEM section and *c* is the third axis, set equal to the longer of the two measured axes (consistent with the oblate-spheroid chloroplast morphology observed by CLSM). Total gyrobody membrane area was estimated from the SPIRE-fitted model as 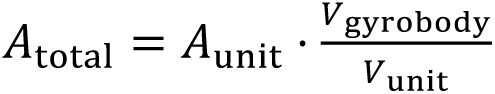 where *A*_unit_ is the total membrane area within a unit cell (as per the formulae above and consistent with estimates obtained by SPIRE) and *V*_unit_ = *a*^3^ the unit-cell volume (with *a* fitted by SPIRE). The ratio 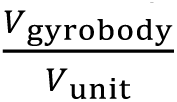 gives the number of unit cells contained in the gyrobody.

The membrane area of an average granum stack was calculated from the GRANA-derived parameters by approximating the stack as a cylinder composed of *N*disk-shaped thylakoids, each contributing two membrane faces: 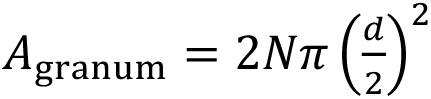 where *d* is the granum diameter and *N* is the number of thylakoids in the stack. Grana stack diameter, height, and the resulting average-granum membrane area were obtained from *stn7-1* samples collected after 16 h of darkness.

### PAM Chlorophyll fluorescence imaging

Chlorophyll fluorescence was measured using an IMAGING-PAM M-Series MAXI system (Walz). 16 h darkness samples were measured. Minimal fluorescence (F₀) was recorded with blue measuring light at 0.5 μmol photons m⁻² s⁻¹. Maximum fluorescence (F_M_) was induced with a 0.84 s saturating blue pulse of 2700 μmol photons m⁻² s⁻¹. After 60 s dark relaxation, plants were exposed to red actinic light at 35 μmol photons m⁻² s⁻¹ for 200 s, with saturating pulses applied every 20 s. Whole plants were used as regions of interest, with fluorescence values collected from multiple spots on leaves. Data were acquired using ImagingWinGigE.

### Thylakoid surface charge

Thylakoid surface charge was assayed using 9-aminoacridine fluorescence quenching. Thylakoids equivalent to 100 μg chlorophyll were washed through three cycles in 0.5 ml buffer E containing 10 mM HEPES-NaOH, pH 7.5, 0.1 M sorbitol, and 1 mM EDTA, with centrifugation at 10,000 × *g* for 3 min at 4°C. Membranes were resuspended to 1.5 mg ml⁻¹ chlorophyll in buffer E. Fluorescence was measured in a stirred 1 cm quartz cuvette on a Shimadzu RF5301-PC fluorometer using excitation at 340 nm and emission at 455 nm. Excitation and emission slit widths were 5 nm and 1 nm, respectively. Measurements were performed in buffer F containing 1 mM HEPES-NaOH, pH 7.6, 0.1 M sorbitol, and 1 mM EDTA. 9-aminoacridine and DCMU were added to final concentrations of 20 μM each. Thylakoids were added to 10 μg ml⁻¹ chlorophyll concentration. After fluorescence reached a plateau, 3 μl of 50 mM EDTA was added, followed by MgCl₂ to the final concentration of 20 mM. Fluorescence traces were normalized to the maximal fluorescence signal (F_max_) in the presence of thylakoids. F/F_max_ values were compared between washed thylakoid samples. For details see (*79*).

### Laurdan membrane fluidity analysis

Thylakoid membranes were diluted to 2.5 μg ml⁻¹ chlorophyll in buffer containing 20 mM HEPES-NaOH, pH 7.5, and 330 mM sorbitol, and equilibrated for 10 min at room temperature. Laurdan was added to 1 μM, and samples were incubated for 30 min at room temperature. Fluorescence steady-state spectra were recorded on a Shimadzu RF-5301PC fluorometer with excitation at 390 nm, excitation slit width of 10 nm, emission from 400 to 600 nm, and emission slit width of 5 nm. Generalized polarization was calculated as 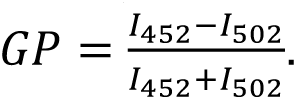.

### HPLC analysis of carotenoids

Carotenoids were extracted from thylakoid membranes or membrane fractions equivalent to 100 μg chlorophyll. Samples were mixed with 500 μl acetone:ethyl acetate (3:2, v/v), vortexed for 15 s at 4°C, mixed with 400 μl cold water, and inverted five times. Samples were centrifuged at 800 × *g* for 5 min at room temperature, and the upper ethyl acetate phase was collected. The lower phase was re-extracted with 200 μl ethyl acetate. Combined extracts were evaporated under argon, resuspended in methanol:hexane:isopropanol (6:3:1, v/v/v), and stored at −80°C until analysis.

HPLC was performed on a Shimadzu Prominence system equipped with photodiode-array detection. Samples were separated on an Atlantis C18 column (4.6 × 250 mm, 5 μm, 100 Å) with an Atlantis dC18 guard column (3.9 × 5 mm, 5 μm). The injection volume was 5 μl. Separation was performed at 1 ml min⁻¹ using a gradient of ethyl acetate in acetonitrile:water:triethylamine (9:1:0.01, v/v/v): 0% ethyl acetate for 2 min, 0 to 66.7% from 2 to 32 min, and 66.7 to 100% from 32 to 37 min. Peaks were integrated at 456 nm (*80*). Pigments were identified by retention time and PDA spectra as described in (*80*).

### High Performance Thin-Layer chromatography (HPTLC) of thylakoid lipids

Polar lipids were extracted from thylakoid membranes or membrane fractions equivalent to 100 μg chlorophyll. Samples were mixed with 5 ml chloroform:methanol (2:1, v/v), shaken for 5 min at 4°C, and centrifuged at 2000 × *g* for 15 min at 4°C. Pellets were re-extracted with 5 ml methanol:chloroform:water (2:1:0.8, v/v/v). Combined extracts were mixed with 2 ml chloroform and 2 ml 0.2 M KCl for 15 min. The lower organic phase was collected, evaporated at 50°C and 200 rpm on a Laborota 4000 rotary evaporator, and resuspended in 1 ml chloroform:methanol (1:1, v/v).

Lipids were separated on HP-TLC Silica gel 60 plates (Supelco) coated with ammonium sulfate and activated at 100°C for 1 h. Samples were applied as 5 mm bands using a CAMAG Linomat 5, and plates were developed in acetone:toluene:water (91:30:7.2, v/v/v). Lipids were visualized with 0.1% (w/v) primuline in 80% (v/v) acetone, imaged at 366 nm and in visible light using a TLC Visualizer 2, and quantified in visionCATS software. MGDG, DGDG, SQDG, and PG standards were from Avanti Polar Lipids.

### Abscisic acid (ABA) quantification

ABA was quantified using the ABA ELISA kit (Agdia). 1.5 g frozen leaf tissue was ground in liquid nitrogen and extracted overnight at 4°C in the dark with 15 ml 80% (v/v) methanol containing 0.01% (w/v) butylated hydroxytoluene and 0.05% (w/v) citric acid monohydrate. Extracts were filtered, evaporated, dissolved in 3.5 ml 10% (v/v) methanol in kit TBS buffer, and diluted fourfold in TBS.

ELISA reactions were performed in duplicate wells with 100 μl standards or samples and 100 μl tracer for 3 h at 4°C. Plates were washed three times with PBST, developed with 200 μl substrate for 60 min at 37°C, and read at 405 nm. ABA concentrations were calculated using the kit binding and logit transformation procedure. An internal ABA standard was used at 0.8 pmol ml⁻¹. ABA values were normalized to fresh weight.

### SDS-PAGE and immunoblotting

Thylakoid samples equivalent to 1 or 3 μg chlorophyll were denatured for 10 min at 75°C in Laemmli buffer containing 138 mM Tris-HCl, pH 6.8, 6 M urea, 22.2% (v/v) glycerol, 4.3% (w/v) SDS, 150 mM β-mercaptoethanol, and 0.01% (w/v) bromophenol blue. Samples were centrifuged at 3000 × *g* for 2 min at room temperature before loading. Proteins were separated on 1 mm SDS-PAGE gels using a Mini-PROTEAN Tetra system (Bio-Rad) at 15 mA per gel for approximately 1.5 h. Resolving gels contained 420 mM Tris-HCl, pH 9.18, 14% (w/v) acrylamide, 0.1% (w/v) bisacrylamide, 0.1% (w/v) SDS, 12% (w/v) sucrose, 0.05% (v/v) TEMED, and 0.025% (w/v) APS. Stacking gels contained 54 mM Tris-HCl, pH 6.1, 5% (w/v) acrylamide, 0.08% (w/v) bisacrylamide, 0.1% (w/v) SDS, 0.25% (v/v) TEMED, and 0.05% (w/v) APS. The cathode running buffer contained 12.5 mM Tris-glycine, pH 8.4, 96 mM glycine, and 0.05% (w/v) SDS.

For immunoblotting, proteins were transferred to methanol-activated 0.2 μm PVDF membranes (Bio-Rad), using a wet Trans-Blot system at 100 V for 45 min at room temperature with cooling and stirring. The transfer buffer contained 25 mM Tris-HCl, pH 8.3, 192 mM glycine, and 10% methanol. Membranes were blocked for 1 h at room temperature or 16 h at 4°C in TBS containing 5% (w/v) milk, or 5% (w/v) BSA for phosphoprotein detection. Primary antibodies (Agrisera) against Lhcb2 (AS01 003, 1:2000), Lhcb2-P (AS13 2705, 1:10 000), Lhcb5 (AS01 009, 1:2000), Lhca2 (AS01 006, 1:5000), PetA (AS20 4377, 1:1000), D1 (AS05 084, 1:10 000), D1-P (AS13 2669, 1:10 000), CP43 (AS11 1787, 1:3000), CURT1A (AS08 316, 1:1000), were used. HRP-conjugated goat anti-rabbit secondary antibody (AS09 602) was used at 1:5000. Signals were detected with Clarity Western ECL (Bio-Rad) and quantified in ImageLab 6.0.1 (Bio-Rad) using Col-0 16 hD signal intensity as a reference.

### Pro-Q Diamond and SYPRO Ruby Staining

For total protein staining, SDS-PAGE gels were fixed twice for 30 min in 50% (v/v) methanol and 7% (v/v) acetic acid at room temperature with gentle agitation, then stained overnight at 4°C with SYPRO Ruby protein gel stain (Invitrogen) protected from light. Gels were washed once for 30 min in 10% (v/v) methanol and 7% (v/v) acetic acid, rinsed twice for 5 min in water, and imaged on a ChemiDoc MP using the SYPRO Ruby acquisition program.

For phosphoprotein staining, SDS-PAGE gels were fixed overnight at 4°C in 50% (v/v) methanol and 10% (v/v) acetic acid with gentle agitation. Gels were washed three times for 10 min in water, stained with Pro-Q Diamond phosphoprotein stain (Invitrogen) for 75 min at room temperature, destained three times for 30 min in 50 mM sodium acetate, pH 4.0, containing 20% (v/v) acetonitrile, and washed four times for 5 min in water. Gels were protected from light after staining and imaged on a ChemiDoc MP using the Pro-Q Diamond acquisition program.

After Pro-Q imaging, gels were stained overnight at 4°C with SYPRO Ruby (Invitrogen) washed once in 10% (v/v) methanol and 7% (v/v) acetic acid, washed twice in water, and imaged on a ChemiDoc MP using the SYPRO Ruby acquisition program.

### Low-temperature (77 K) fluorescence spectroscopy

Thylakoids were diluted to 10 μg ml⁻¹ total chlorophyll in buffer G containing 20 mM HEPES-NaOH, pH 7.5, 330 mM sorbitol, 15 mM NaCl, and 4 mM MgCl₂. Samples were frozen in liquid nitrogen and measured at 77 K using a modified Shimadzu RF5301-PC spectrofluorometer with a Teflon cuvette mounted in a quartz holder connected by optical fibers. Fluorescence emission was recorded from 600 to 800 nm at 1 nm intervals with excitation at 412, 440, or 470 nm. Excitation and emission slit widths were 5 nm, and an LP600 emission filter was used. Buffer spectra were subtracted, and spectra were normalized to equal area over the entire collected wavelength range.

### Circular Dichroism (CD) spectroscopy

CD spectra were recorded from thylakoids at 10 μg ml⁻¹ total chlorophyll in buffer G using a 1 cm quartz cuvette and a Chirascan-plus spectropolarimeter (Applied Photophysics). Spectra were acquired from 350 to 800 nm at 1 nm intervals and 1 nm bandwidth with simultaneous absorbance recording and were normalized to the chlorophyll Qy absorption band.

### FTIR Spectroscopy

Thylakoids equivalent to 50 μg chlorophyll were washed four times for 5 min at 10°C in D₂O-based buffer C and collected by centrifugation at 7200 × *g* for 5 min at 10°C. The final pellet was resuspended in D₂O-based buffer C to 5 mg ml⁻¹ chlorophyll. A 5 μl aliquot was deposited on a diamond/ZnSe ATR crystal and partially dehydrated under nitrogen to form a thin film. Spectra were acquired in the dark under nitrogen on a Shimadzu IRAffinity-1S spectrometer equipped with a MIRacle ATR single-reflection accessory. Ten series of 20 interferograms were acquired, averaged, and Fourier-transformed. Data were restricted to the range of 1800-1580 cm⁻¹, and a linear baseline was subtracted to the minima in the selected range, which includes the C=O stretching vibration of the ester group of fatty acids/glycerol esters (1790–1590 cm^−1^) and the Amide I (1710-1585 cm⁻¹) regions. The spectra were then normalized to equal area under the curve.

### Fluorescence Lifetime Correlation Spectroscopy

FLCS measurements were performed on intact isolated chloroplasts at 25°C, maintained in a custom-built climate chamber, using a Nikon Eclipse TE2000U microscope (Nikon) coupled to a PicoHarp 300 system (PicoQuant) and controlled by SymPhoTime 64 software (PicoQuant) integrated with NIS-Elements software (Nikon). Samples were immobilized on poly-L-lysine-coated glass. Excitation was provided by a 485 nm pulsed diode laser at 1 to 5 μW, and fluorescence was collected through a 488 nm long-pass filter using a 60× water-immersion objective (NA 1.2) and SPAD (Single Photon Avalanche Diode) detectors with a preceding filter (525/50 nm). The instrument was calibrated with 50 nM rhodamine 110 in 20 mM HEPES-NaOH buffer (pH 7.5) containing 330 mM sorbitol, 15 mM NaCl, 4 mM MgCl₂, and 10 mM NaF, following the procedure described in (81).

For FLCS measurements, thylakoids were incubated in the same buffer with or without 50 nM rhodamine 110. Time-resolved fluorescence fluctuation signals were acquired from manually selected thylakoid regions exhibiting cubic or lamellar membrane structure.

Because thylakoids exhibit strong autofluorescence in the visible range, photon filtering based on fluorescence lifetime was applied to remove this residual signal. The autofluorescence decay of thylakoids without added tracer was first acquired under 485 nm pulsed excitation and analyzed with the TCSPC module of SymPhoTime 64, yielding a characteristic lifetime of ∼0.2 ns (fig. S13A). Autocorrelation of the same signal showed no diffusion-like pattern, confirming the absence of mobile autofluorescent species in the thylakoid structure (fig. S13B). For rhodamine 110 measurements, the autofluorescence lifetime decay was then used as a filter in the FLCS module of SymPhoTime 64 to discard photons most likely originating from the thylakoid itself, after which the remaining photons were correlated according to the standard FCS procedure. Without this filtering, autocorrelation curves were dominated by autofluorescence-derived photons rather than the tracer (fig. S13C–E).

Filtered autocorrelation curves were fitted using FcsIT software (*82*) with a two-component free diffusion model including a triplet-state term (Eq. S1, fig. S13F), accounting for the freely diffusing dye (*τ_D_*_1_) and an additional component attributed to unspecific binding to membranes or proteins (*τ_D_*_2_) (*83*).

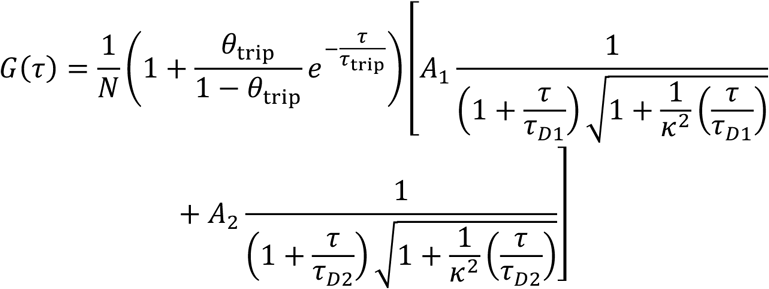

The structural parameter (κ, focal-volume aspect ratio) was determined during calibration and fixed during fitting. Following (*83*), *τ_D_*_1_ was used in downstream analysis, and diffusion coefficients were calculated as 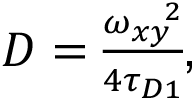, where *ω_xy_* is the focal-volume radius determined during calibration (*81*).

## Acknowledgments

We are grateful to I. Finkemeier for providing seeds of the Arabidopsis thaliana *gnat2-1* mutant and to M. Pribil for sharing *oe*STN7 seeds. We warmly thank M. Hapka for proposing the term “gyrobody,” which gave a name to the central finding of this work. We thank J. Kozioł-Lipińska, B. Bieniak, and T. Góral for their excellent technical assistance. Room-temperature TEM studies were performed in the Laboratory of Electron Microscopy of the Nencki Institute of Experimental Biology, supported by a project financed by the Polish Minister of Education and Science under contract no. 2022/WK/05 (Polish Euro-BioImaging Node “Advanced Light Microscopy Node Poland”). FTIR experiments were conducted in the Laboratory of Biophysics, Chemistry and Molecular Biology at the Faculty of Physics, University of Warsaw. Cryo TEM analysis of isolated membranes was performed at Cryomicroscopy and Electron Diffraction Core Facility, Centre of New Technologies, University of Warsaw.

## Author contributions

MB performed SPIRE analysis, 3D modeling from TEM and CLSM data, spatial parameter computations, carotenoid analysis, developed theoretical models, prepared cryo-ET samples, designed and prepared figures and videos, and wrote the Materials and Methods section. AW performed FTIR, circular dichroism (CD), and 77 K chlorophyll fluorescence measurements, cryo-EM and HPF-FS of isolated membranes, and membrane fractionation with SDS-PAGE analysis and Western blot analysis. WW performed cryo-ET analysis, segmentation, and modeling, and prepared figures. AB performed TEM of tissues and isolated membranes, GRANA analysis, manual measurements on TEM micrographs, ABA level assessment, Western blot analysis, and Pro-Q Diamond staining. JW performed HP-TLC of lipids, Western blot analysis, Pro-Q Diamond staining, membrane fractionation, carotenoid analysis, and ABA level assessment. RM performed membrane charge analysis, FLCS analysis, CLSM analysis, and PAM imaging analysis, and prepared figures. FLB performed cryo-ET analysis and segmentation. ZB performed Laurdan fluorescence measurements and statistical analysis, and prepared figures. KK performed FLCS analysis and provided the FLCS instrumentation. GEST supervised the geometrical analysis and advised on the mathematical sections of the manuscript. BDE provided access to cryo-ET infrastructure and supervised the cryo-ET analysis. ŁK conceived the study, secured funding, performed TEM analysis of tissues, CLSM analysis, and membrane charge analysis, designed figures, supervised the project, and wrote the manuscript. All authors reviewed and edited the manuscript.

## Data, code, and materials availability

Raw data, images, and scripts associated with the figures and analysis are available at https://danebadawcze.uw.edu.pl, https://doi.org/10.58132/VNNZV4.

Cryo-electron tomography raw data have been submitted to the Electron Microscopy Public Image Archive (EMPIAR) and are currently under deposition. Cryo-ET data containing raw, denoised, and segmented tomograms used to generate figures in this manuscript have been deposited in the Electron Microscopy Data Bank (EMDB) under accession codes EMD-58572, EMD-58573, and EMD-58574. The EMPIAR accession number will be provided upon completion of the deposition process and included in the final published version of the manuscript

## Supplementary Materials

Figs. S1 to S13

Video S1

---

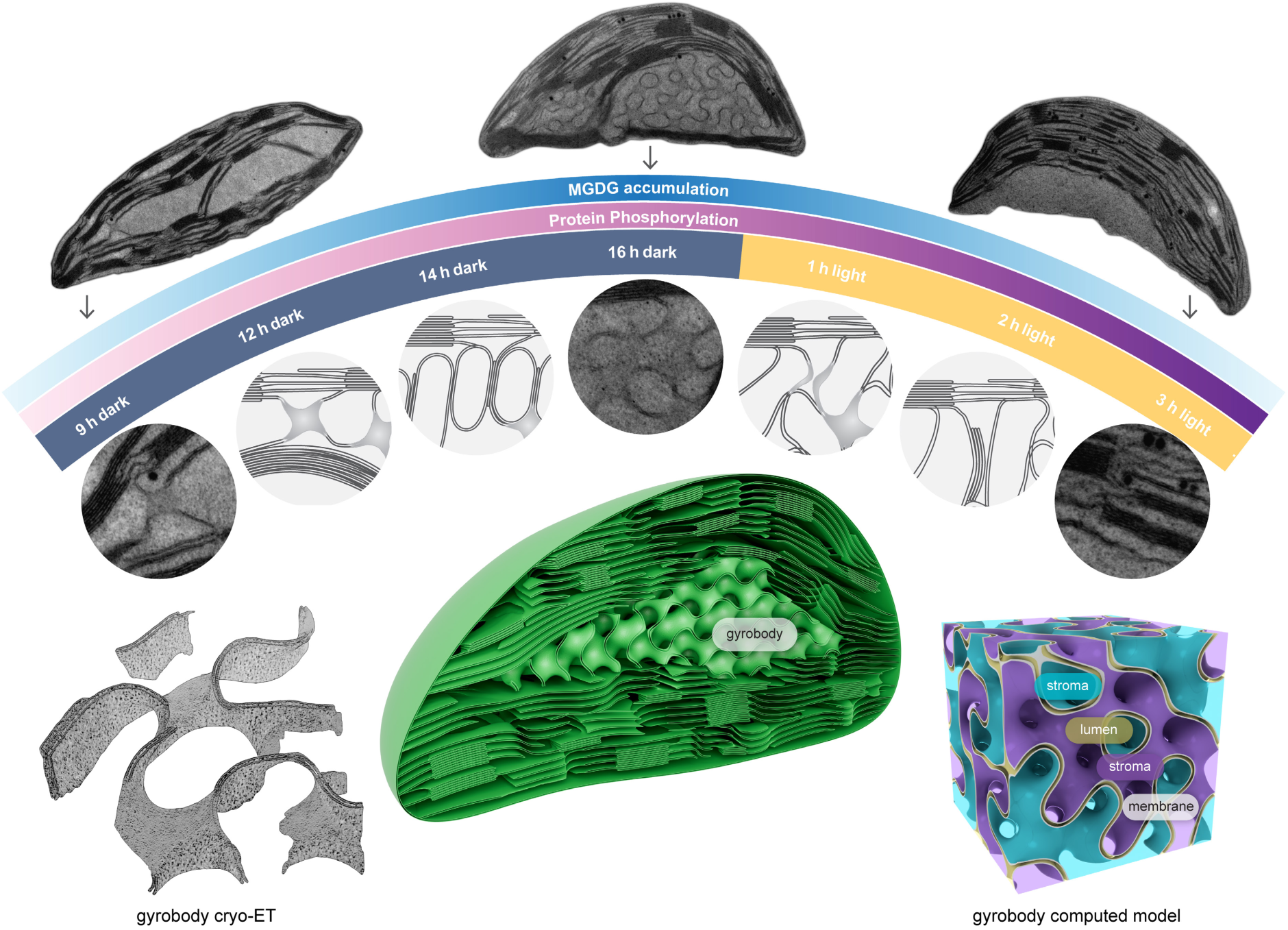
Summary figure. Mature plant chloroplasts adopt a gyroid cubic membrane configuration. A convoluted membrane form, termed gyrobody, forms reversibly during the day/night cycle through a two-stage molecular process: reduced thylakoid phosphorylation lowers membrane surface charge and triggers grana disassembly, after which monogalactosyldiacylglycerol (MGDG) accumulation facilitates membrane folding into a gyroid form. With a ∼500 nm unit cell, the gyrobody occupies a substantial fraction of the chloroplast volume, accommodates three distinct aqueous channels, and reorganizes the distribution of photosystems.

