## Supplementary figures for "Mature plant chloroplasts form reversible gyroid cubic membranes"

for

**Vid. S1. The mature chloroplast cubic membrane is based on a gyroid-type minimal surface that divides space into three interpenetrating but distinct aqueous channels.** The video shows a close-up of a representative gyrobody from an *aba1-6* mutant, followed by sequential steps: projection matching with the warping step, TEM slice reconstruction (coral), and the resulting reconstruction of a larger gyroid region with visualization of the three interpenetrating aqueous channels.

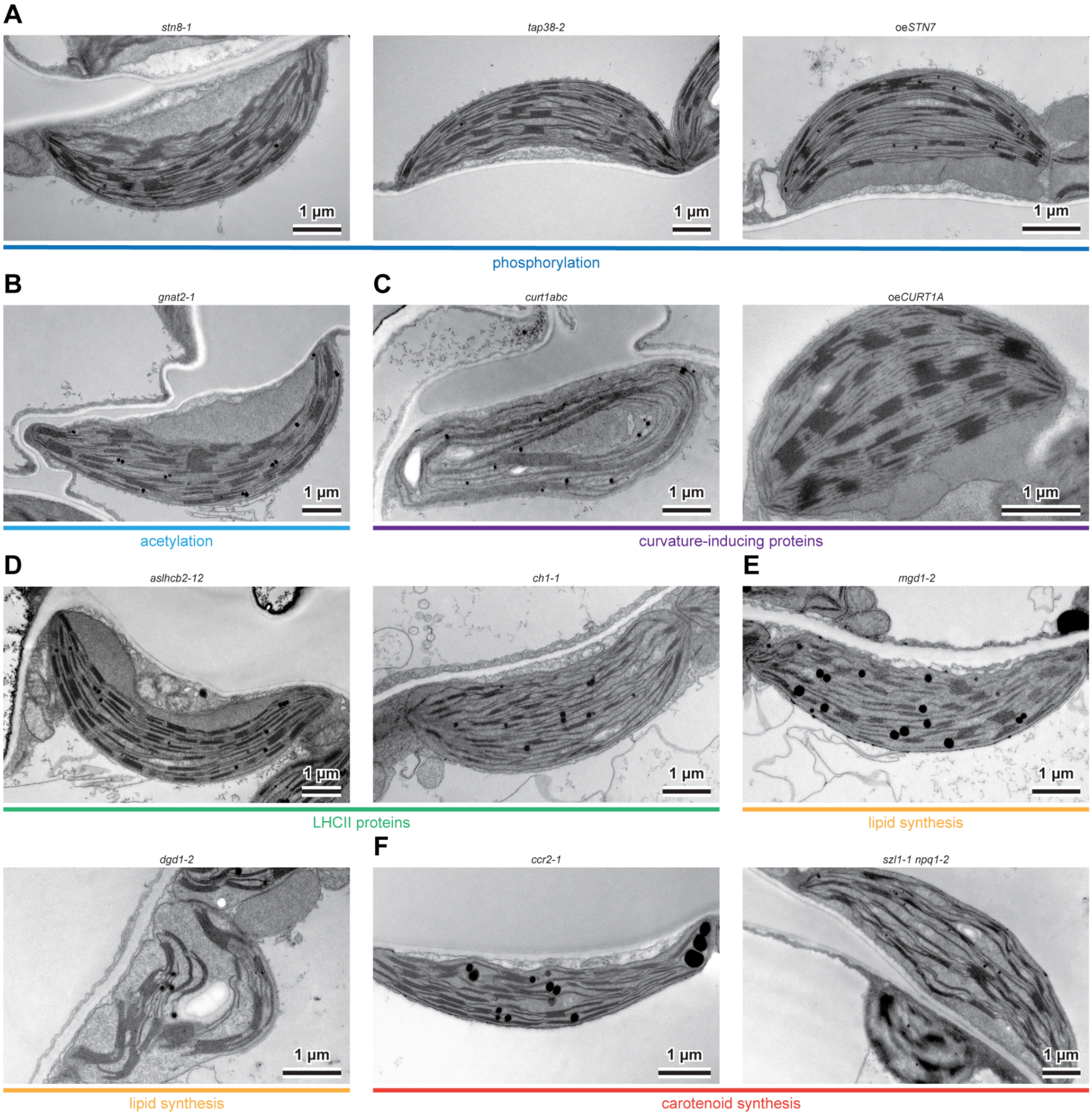


**Fig. S1. Thylakoid networks of mature chloroplasts retain lamellar configuration in multiple mutants with altered membrane composition.**

(A–F) Representative TEM images of mature Arabidopsis chloroplasts. Samples were collected at the end of the night period (16 hD) from fully mature leaves of 6–8 week-old plants. (A) Plants with disturbed thylakoid protein phosphorylation (*stn8-1*, *tap38-2*, oe*STN7*). (B) Plant depleted in thylakoid protein acetylation (*gnat2-1*). (C) Plants with disturbed CURT protein levels (*curt1abc*, oe*CURT1A*). (D) Plants with disturbed LHCII protein levels (*aslhcb2-12*, *ch1-1*). (E) Plants with disturbed thylakoid galactolipid content (*mgd1-2*, *dgd1-2*). (F) Plants with disturbed carotenoid composition (*ccr2-1*, *szl1-1 npq1-2*). Data are representative of 2–3 biological replicates.

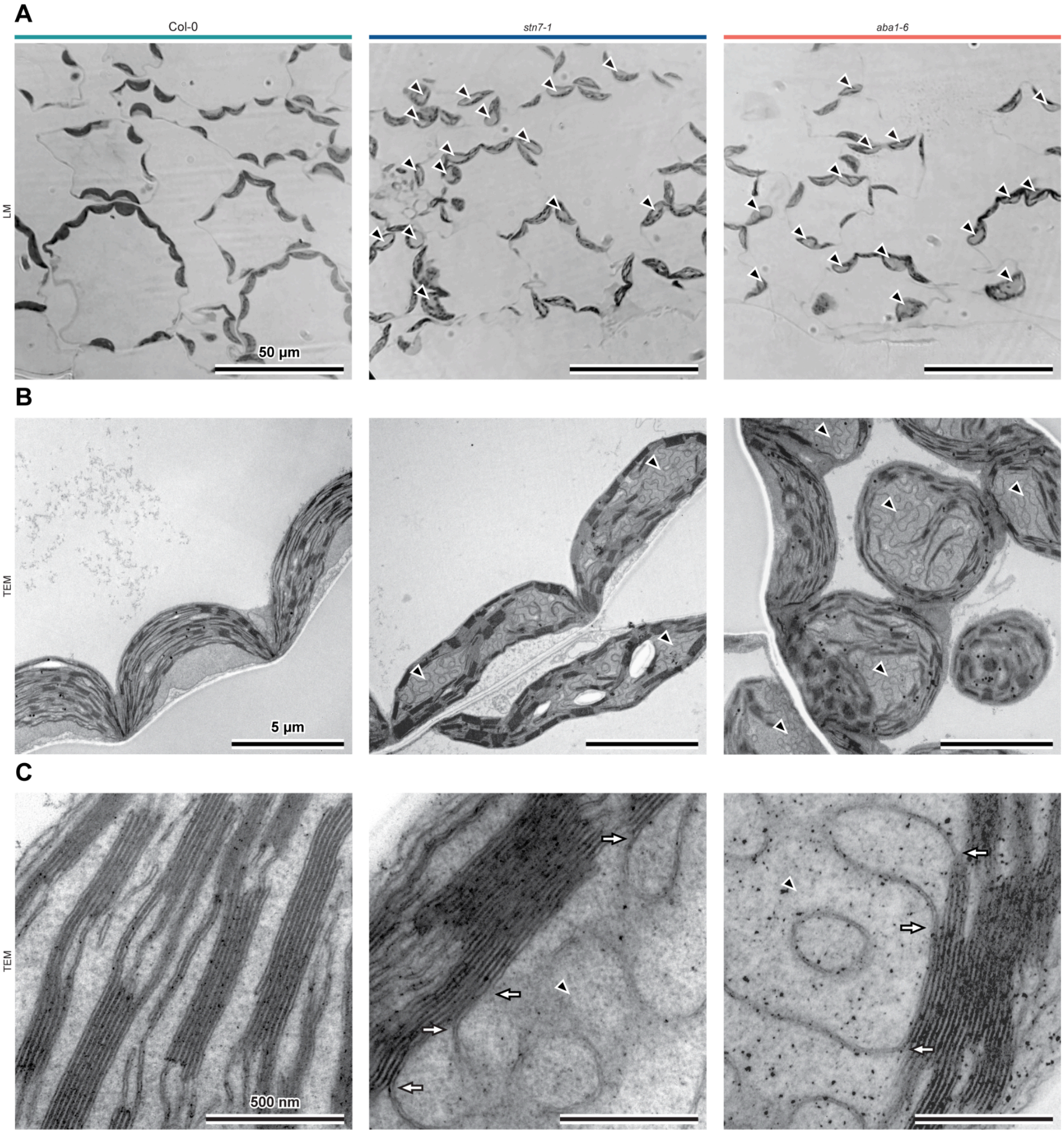


**Fig. S2. Gyrobodies directly connect to grana stacks and are present in the majority of 16 hD chloroplast cross-sections of *stn7-1* and *aba1-6*.** (A) Semithin sections of Epon-embedded samples stained with toluidine blue (shown in black and white) imaged by bright-field light microscopy (LM). Black arrowheads indicate gyrobodies. Scale bars: 50 µm. (B) TEM images of mesophyll cells showing chloroplasts with fully lamellar thylakoids (Col-0) and chloroplasts additionally containing gyrobodies (black arrowheads). Scale bars: 5 µm. (C) TEM close-ups of the thylakoid network showing direct connections (white arrows) between gyrobodies (black arrowheads) and grana stacks. Scale bars: 500 nm. TEM data are representative of at least three biological replicates; LM data are from two biological replicates.

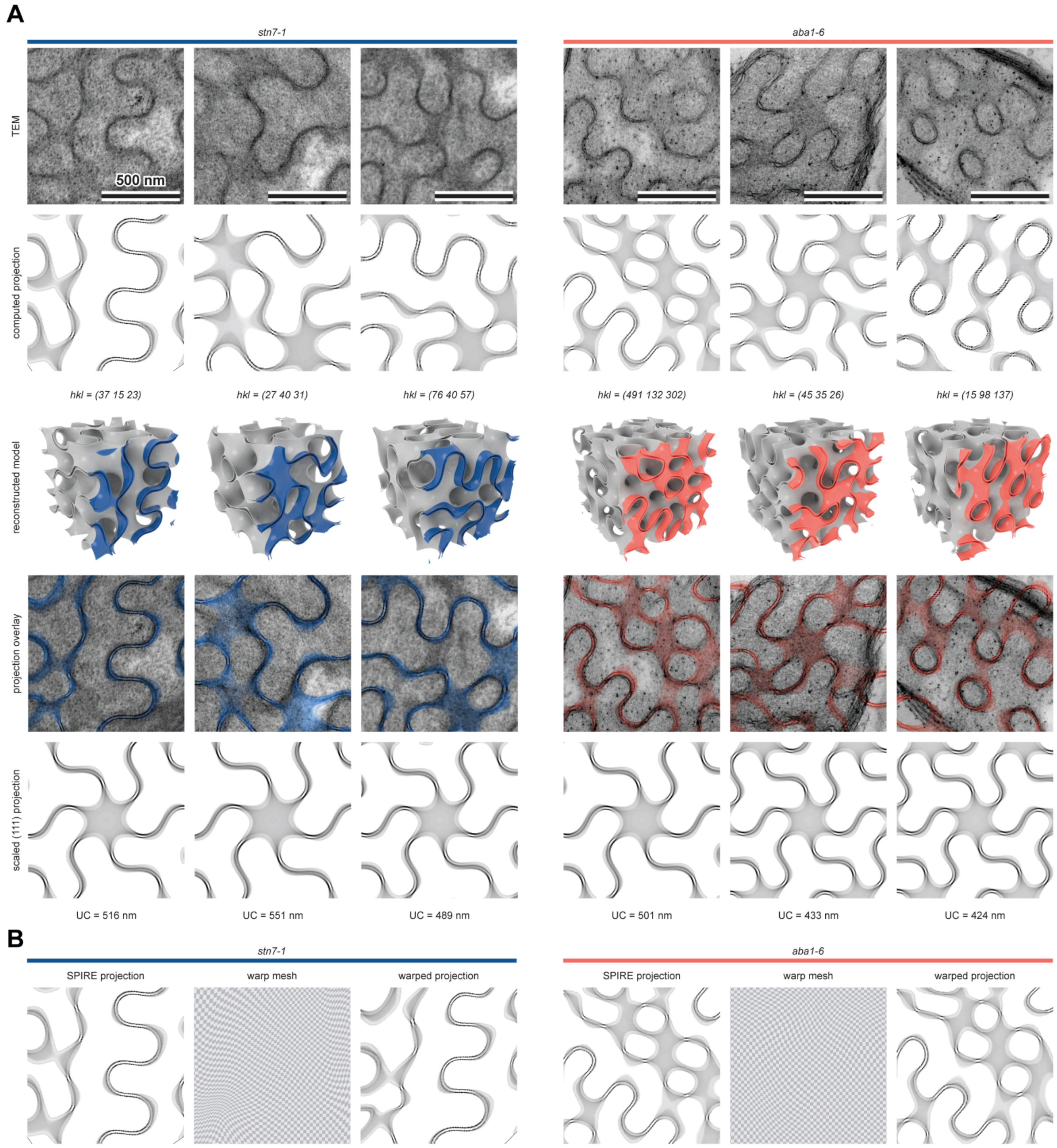


**Fig. S3. Cubic membranes of *stn7-1* and *aba1-6* Arabidopsis mutants are based on a gyroid-type minimal surface.** (A) TEM micrographs of cubic membrane regions selected for template matching analysis (top row). Computed projections were generated interactively using SPIRE (second row), and 3D gyroid models were reconstructed from these projections (third row); colored regions represent the 70 nm TEM specimen thickness. Precise alignment of the computed projections to the TEM images required slight warping (fourth row; see (B) for details), likely reflecting either sample deformation during imaging or the dynamic liquid-crystalline character of large-scale thylakoid gyroids. Unit cell (UC) sizes of the analyzed gyroids range from 424 to 551 nm and vary between genotypes and biological replicates, indicating the dynamic nature of the structure. (B) SPIRE-generated projections with the warp meshes required to align each computed projection with its corresponding TEM image; two representative examples are shown. Data are from three biological replicates.

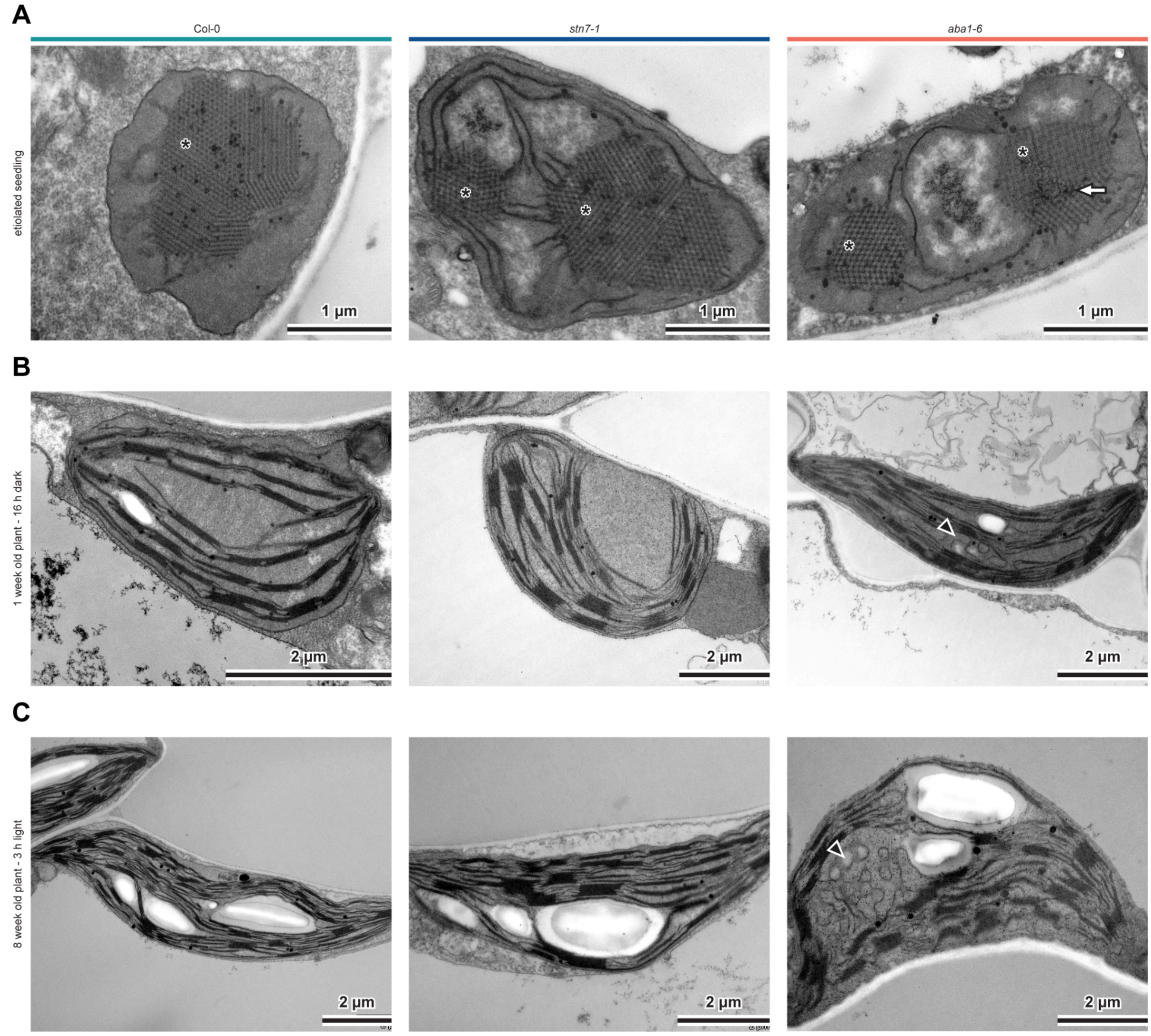


**Fig. S4. Chloroplasts of Col-0, *stn7-1*, and *aba1-6* exhibit different membrane configurations across plastid developmental stages and the day/night cycle.**

(A) TEM images of etioplasts in cotyledons of 5-day-old etiolated seedlings. Diamond-type prolamellar bodies (PLBs) are marked with asterisks. The disturbed cubic PLB network in *aba1-6* etioplasts is indicated by a white arrow. (B) TEM images of chloroplasts in young leaves of 1-week-old plants grown under the day/night cycle, sampled at the end of the dark period. Incipient membrane bending suggesting early gyroid formation is visible in *aba1-6* (black arrowhead). (C) TEM images of chloroplasts in mature leaves of 8-week-old plants grown under the day/night cycle, sampled after 3 h of light. The gyrobody in *aba1-6* is marked with a black arrowhead. Bright starch grains are visible in all genotypes, confirming photosynthetic activity.

Micrographs are representative of at least three biological replicates.

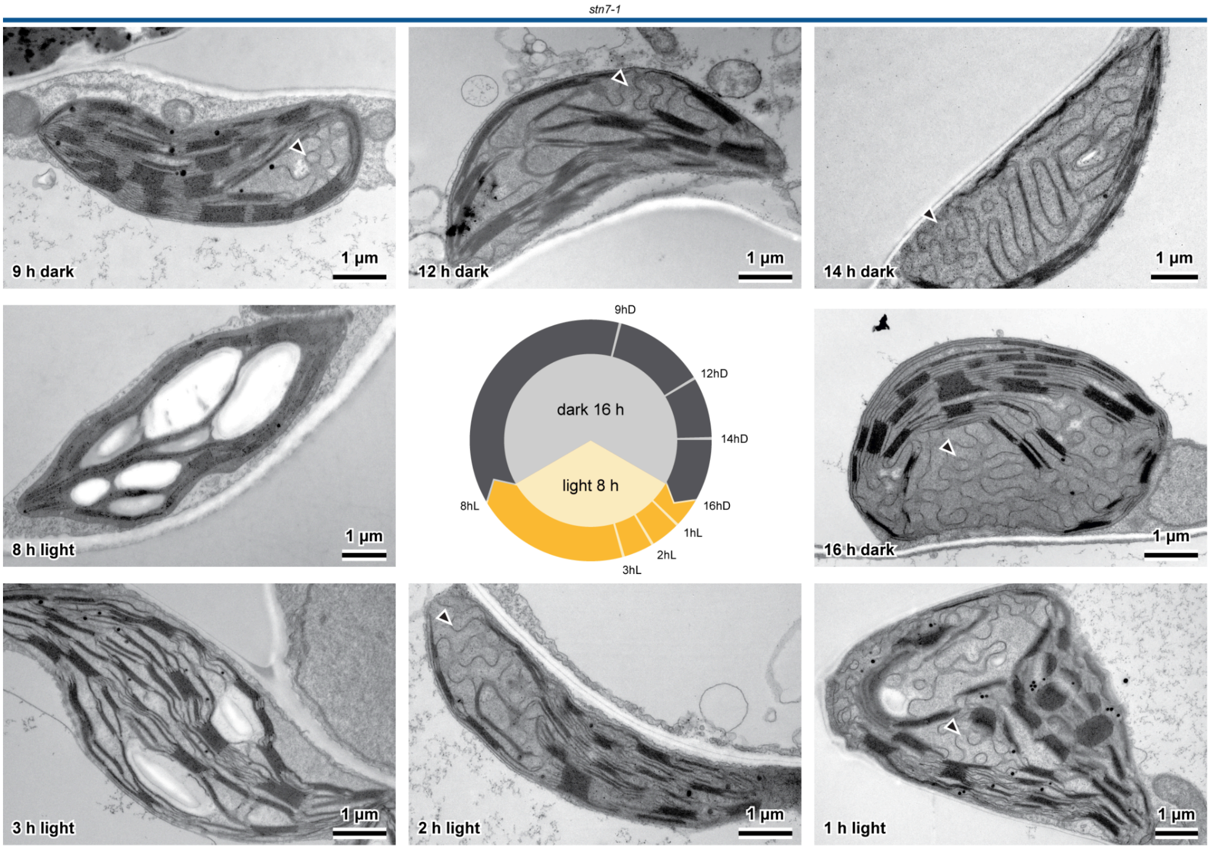


**Fig. S5. Gyrobody formation and disassembly during the day/night cycle in *stn7-1* drives extensive structural reorganization of the thylakoid network.**
Steps of gyrobody formation and disassembly in *stn7-1* plants at selected time points across the day/night cycle. Each stage is shown as a representative TEM image of a whole chloroplast. Gyroid features are marked with black arrowheads. The sampling scheme is shown at the figure center. Scale bars: 1 µm. For close-ups of the membrane transformation, see Fig. 2A.
Data are representative of at least three biological replicates.

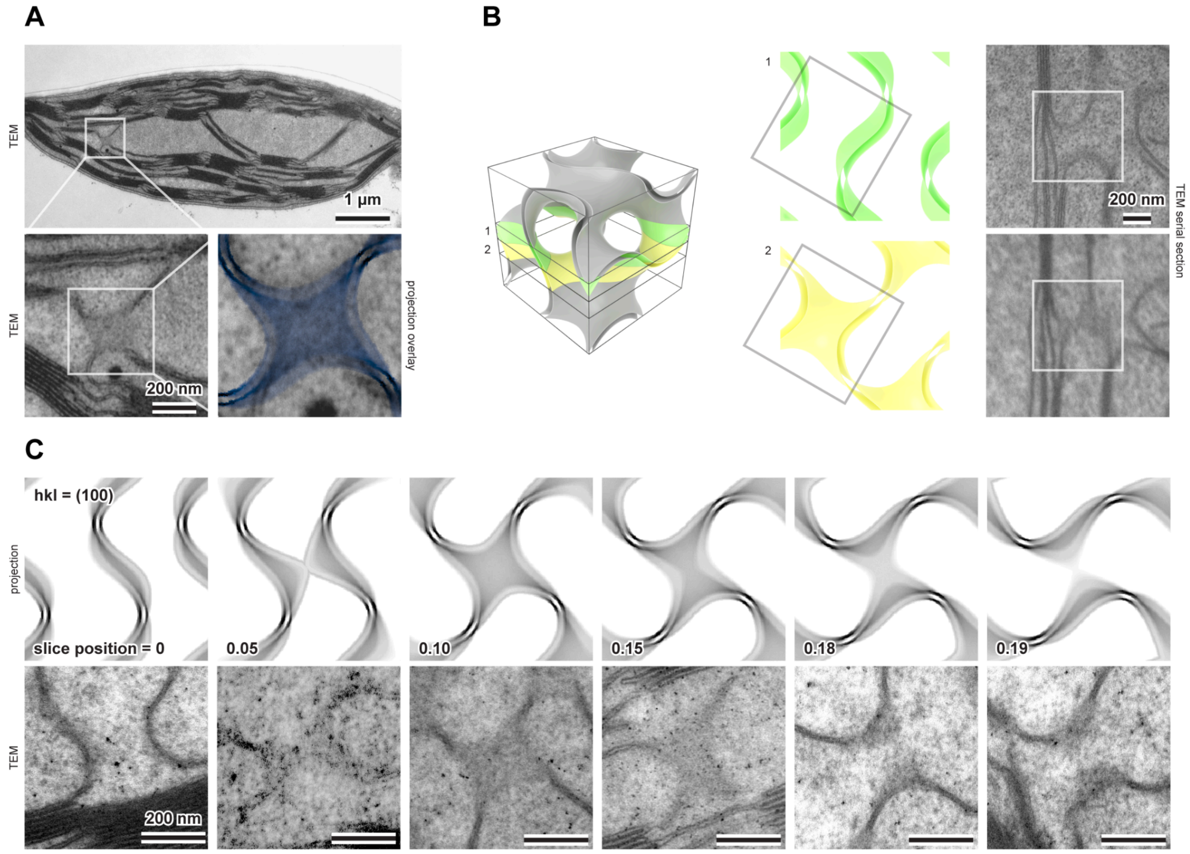


**Fig. S6. Gyrobody formation begins with a single gyroid unit cell, visible in TEM cross-sections in the (100) orientation.** (A) Chloroplast TEM image of an *stn7-1* mutant showing the first sign of gyroid formation: a section through a single gyroid unit cell (UC) in the (100) orientation. (B) Two serial sections through a forming gyroid region showing how the gyroid structure appears different at different z-slice positions. The left side shows a 3D model of a single gyroid UC, with two parallel planes marking the positions of the serial sections (each 70 nm thick, matching the TEM specimen thickness). The right side shows the corresponding TEM images from the same sample region. (C) A selected region of the (100) gyroid projection shown at successive z-slice positions (0–0.19), illustrating how the same structure appears different depending on the depth of the cutting plane. Each SPIRE projection (top row) is paired with a TEM image of similar appearance taken from different samples collected during gyroid formation; these illustrate the range of appearances the same structure can produce in TEM cross-sections. Scale bars: 200 nm, except the top-left image in (A), which is 1 µm.

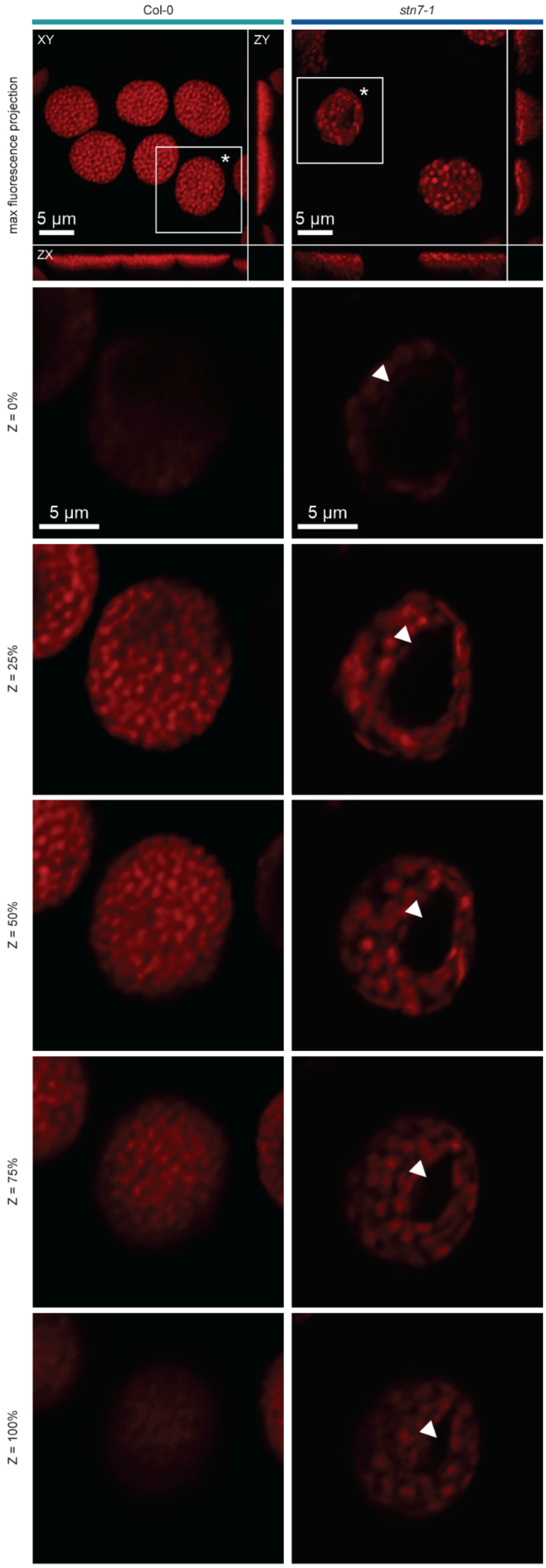


**Fig. S7. The gyrobody appears as a dark non-fluorescent region in CLSM imaging of chloroplast chlorophyll autofluorescence in living mesophyll cells.**

Mesophyll samples for CLSM analysis were collected at the end of the dark period (16 hD) to exclude starch accumulation. (top row) Scanned cell region; the selected chloroplast is marked with an asterisk. (following rows) Optical sections through the same chloroplast in Col-0 and *stn7-1*. Numbers indicate the relative z-position of each optical section, expressed as a percentage of the total axial extent of the chlorophyll fluorescence signal for that chloroplast (0% = top, 100% = bottom). White arrowheads indicate the gyroid region. Data are from at least three biological replicates.

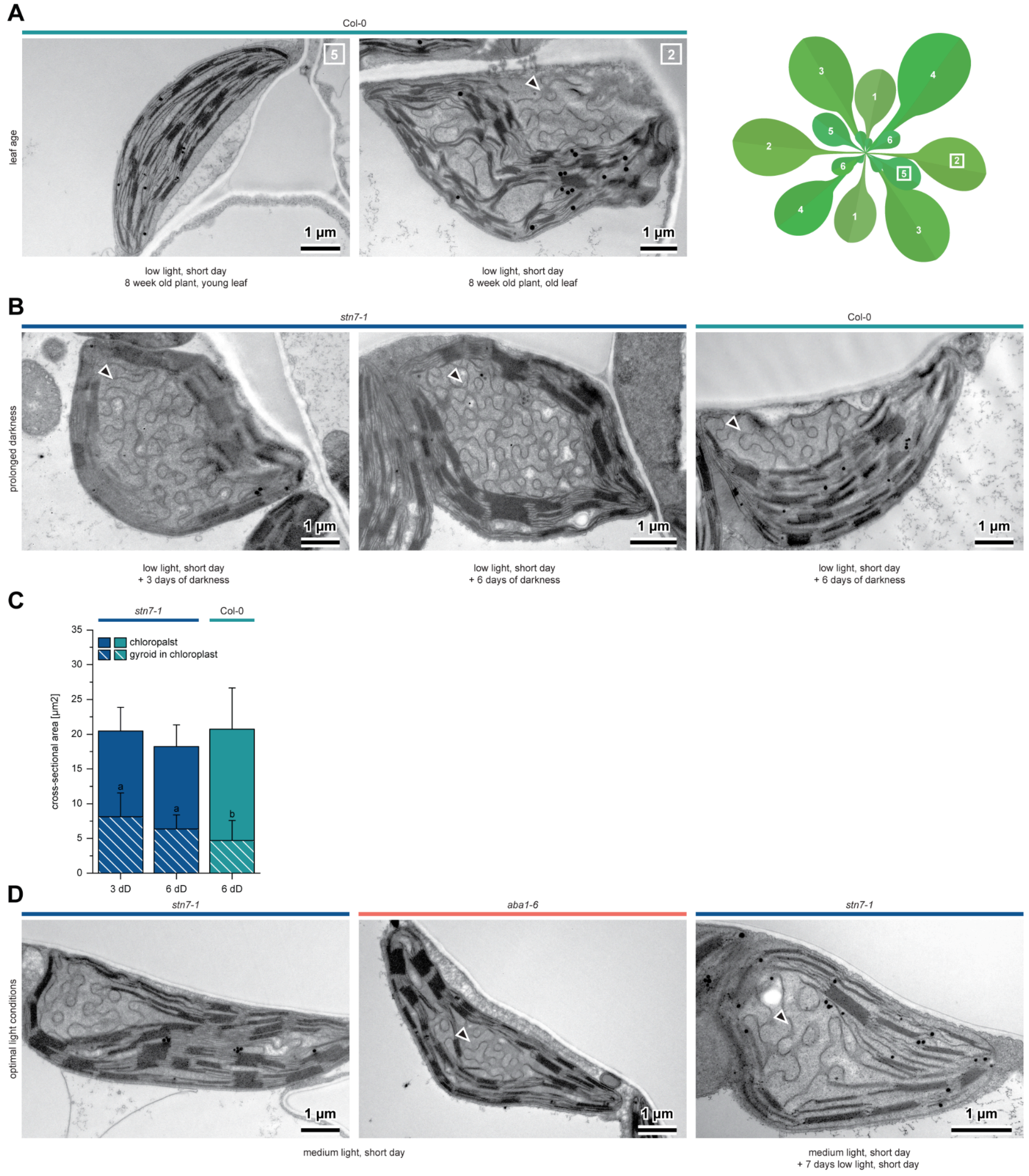


**Fig. S8. Gyrobody size depends on plant growth conditions, and gyrobody presence correlates with leaf age.** (A) TEM images of Col-0 chloroplasts. Samples were collected at the end of the night (16 hD) from 8-week-old plants grown under the day/night cycle, from top (left) or bottom (middle) leaves of the rosette (shown schematically on the right). Leaves from the central part of the rosette were used in all other experiments in this work. Gyrobodies are present in old-leaf chloroplasts (black arrowhead) without typical symptoms of chloroplast senescence. (B) TEM images of chloroplasts from Col-0 and *stn7-1* plants exposed to prolonged darkness (3–6 days) after 8 weeks of growth under the standard day/night cycle in low light (30 µmol photons $m^{-2}$ $s^{-1}$). Gyrobodies were found in both genotypes in middle-rosette leaves (black arrowheads). (C) Cross-sectional area of chloroplasts (solid bars) and of gyrobodies within them (dashed bars) at selected time points of extended darkness in Col-0 and *stn7-1* plants. Error bars indicate SD. Different letters indicate significant differences ($p < 0.05$, Levene's test for variance homogeneity, subsequently, for chloroplast cross-sectional area Welch ANOVA with no post hoc test need and for gyrobodies within chloroplasts one-way ANOVA with Tukey HSD post hoc test). n = [15–22] chloroplasts per condition. (D) TEM images of *stn7-1* and *aba1-6* chloroplasts from plants grown under optimal medium-light conditions (150 µmol photons $m^{-2}$ $s^{-1}$), sampled at the end of the night (left, middle). (right) *stn7-1* chloroplast from a plant grown for 7 weeks in medium light and transferred to low light for the final week; same sampling conditions. Gyroid structures are marked with black arrowheads. Data are representative of at least three biological replicates (A, B) or two biological replicates (D).

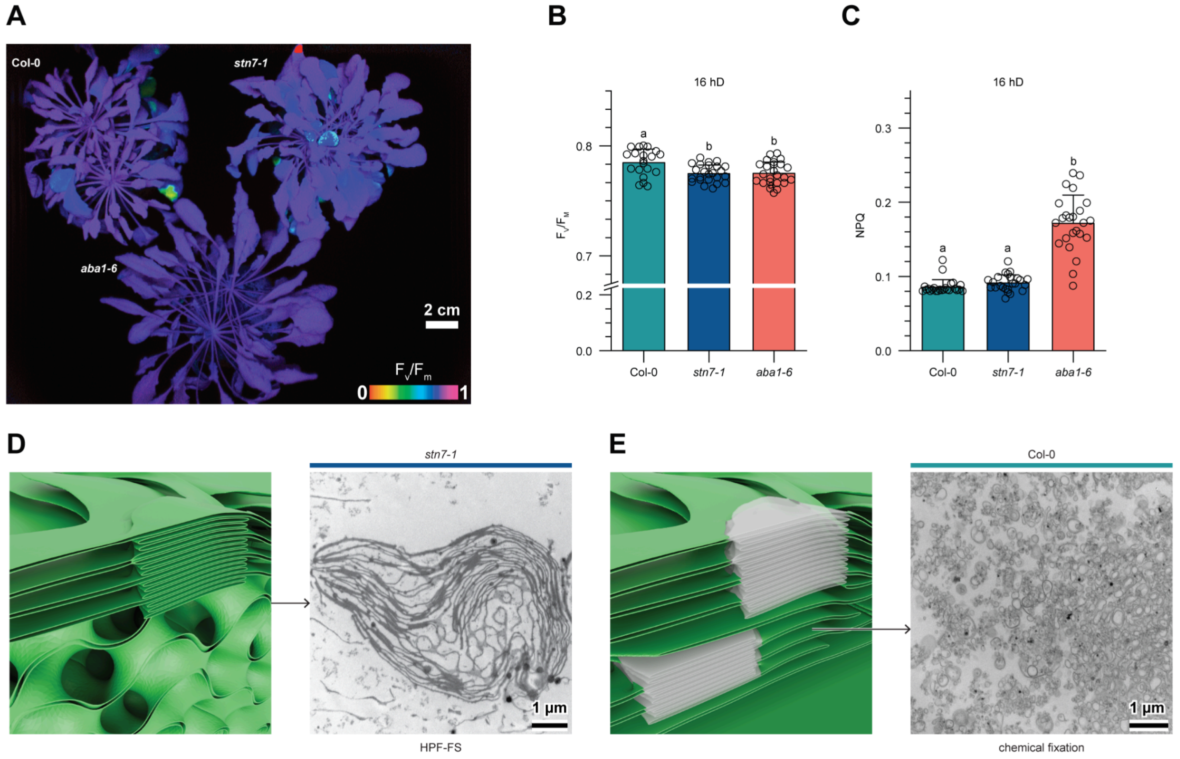


**Fig. S9. Gyrobody presence does not substantially affect chloroplast photosynthetic activity.** (A) Chl *a* fluorescence imaging. Images show rosette morphology and the distribution of the maximum photochemical efficiency of PSII ($F_V/F_M$) in Col-0, *stn7-1*, and *aba1-6* plants at the end of the night period (16 hD). (B–C) $F_V/F_M$ (B) and non-photochemical quenching (NPQ) (C) values for the plants shown in (A). Error bars indicate SD. Different letters indicate significant differences ($p < 0.05$, Levene's test for variance homogeneity, subsequently one-way ANOVA with Tukey HSD post hoc test (B) or Welch ANOVA with Games-Howell post hoc test (C)). n = [20–24] leaf regions per genotype. (D) Isolated thylakoids from *stn7-1* 16 hD samples contain fully developed gyrobodies, visualized by TEM using high-pressure freezing and freeze substitution fixation (HPF-FS), with an accompanying 3D model showing the membrane arrangement at this time point. (E) TEM image of the unstacked thylakoid fraction obtained by digitonin solubilization from Col-0 16 hD samples, paired with a 3D model showing which fractionated region (green) corresponds to the TEM view (black arrow). Panels (D) and (E) extend the data shown in Fig. 3A. Data are representative of three biological replicates.

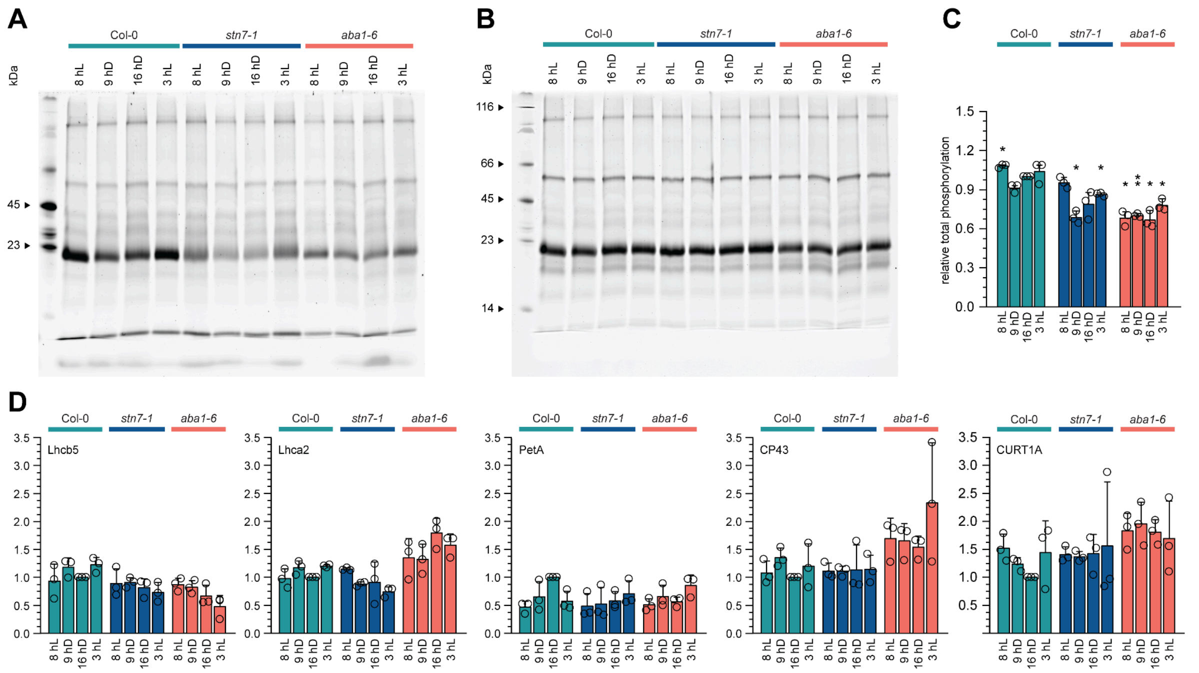


**Fig. S10. Plants with gyrobody at different stages of its formation exhibit decreased thylakoid protein phosphorylation.** (A–B) Phosphoprotein staining (Pro-Q Diamond) of SDS-PAGE-separated thylakoid samples (A) with the corresponding total protein staining (SYPRO Ruby) as a loading control (B), shown for total thylakoid samples of Col-0, *stn7-1*, and *aba1-6* plants across selected time points of the day/night cycle. (C) Relative total phosphorylation calculated from (A). Values are normalized to the Col-0 16 hD sample. Error bars indicate SD. Asterisks indicate significant differences from Col-0 16 hD (*: $p \leq 0.05$, **: $p \leq 0.01$, one-sample *t*-test). n = 3 biological replicates. (D) Densitometric analysis of the remaining immunoblots from Fig. 3B. Values are normalized to the Col-0 16 hD sample. Error bars indicate SD; statistical analysis as in (C). n = 3 biological replicates.

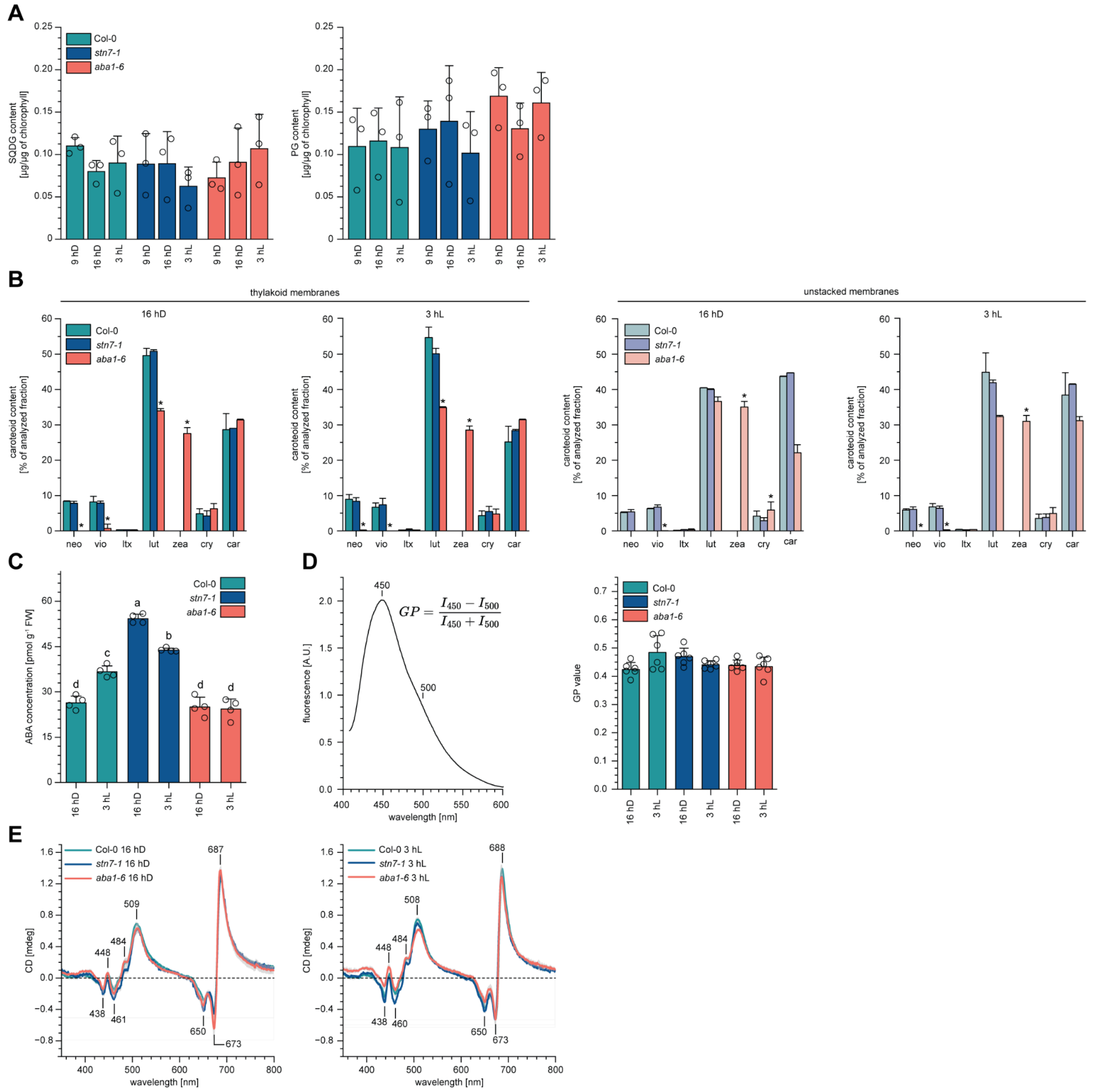


**Fig. S11. Carotenoid and abscisic acid (ABA) content and membrane fluidity are not directly related to gyrobody formation.** (A) Densitometric analysis of SQDG and PG content from the HPTLC analysis in Fig. 3H. Error bars indicate SD. Statistical analysis (Levene's test for variance homogeneity, subsequently one-way ANOVA) revealed no significant differences between samples. n = 3 biological replicates. (B) Relative contribution of individual carotenoids (Neo, Vio, Ltx, Lut, Zea, Cry, Car) to the total carotenoid pool, in total thylakoid (left) and unstacked thylakoid (right) fractions. Error bars indicate SD. Asterisks indicate significant differences ($p < 0.05$, Levene's test for variance homogeneity, subsequently one-way ANOVA with Tukey HSD post hoc test) between genotypes within the same time point for particular carotenoid type. n = 3 biological replicates. (C) ABA concentration per fresh weight in the three studied genotypes at two time points (16 hD, 3 hL). Error bars indicate SD. Different letters indicate significant differences ($p < 0.05$, one-way ANOVA with Tukey HSD post hoc test). n = 4 biological replicates. (D) Representative Laurdan fluorescence emission spectra with the formula for general polarization (GP) calculation (left) and the calculated Laurdan GP values (right) for isolated total thylakoid samples (16 hD and 3 hL) of Col-0, *stn7-1*, and *aba1-6*. Error bars indicate SD; statistical analysis (Levene's test for variance homogeneity subsequently, Welch ANOVA). (E) Full-range CD spectra for the three studied genotypes at two time points (see Fig. 4C–D for the detailed analysis range). Data are from at least three biological replicates.

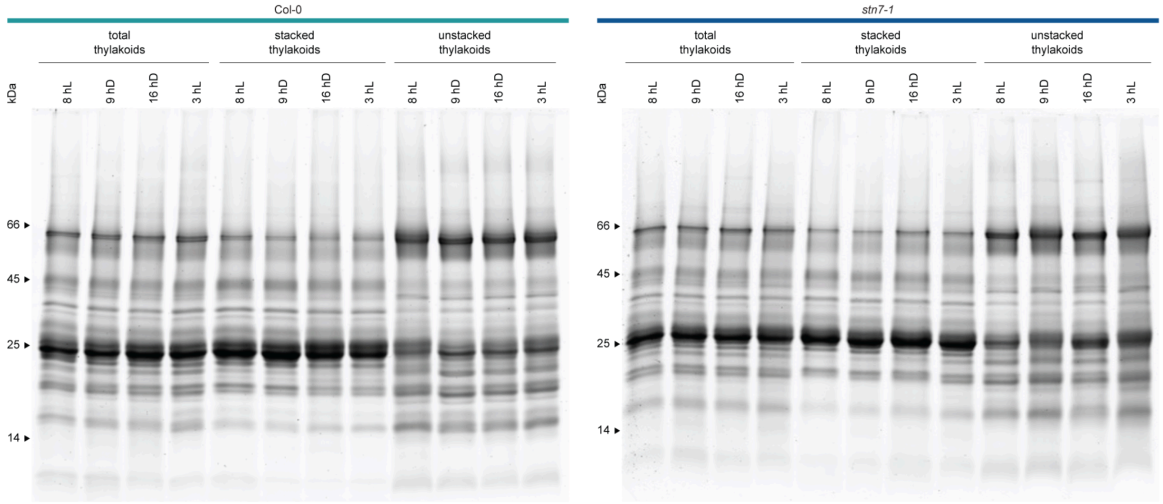


**Fig. S12. LHCII levels are increased in the unstacked thylakoid fraction of *stn7-1* when the gyrobody is fully developed, indicating disrupted lateral segregation of photosynthetic complexes.** SDS-PAGE separation of total, stacked, and unstacked thylakoid fractions from Col-0 and *stn7-1* at selected time points (9 hD, 16 hD, 3 hL, 8 hL) across the day/night cycle. Gels are stained with SYPRO Ruby. Data are from 1 biological replicate.

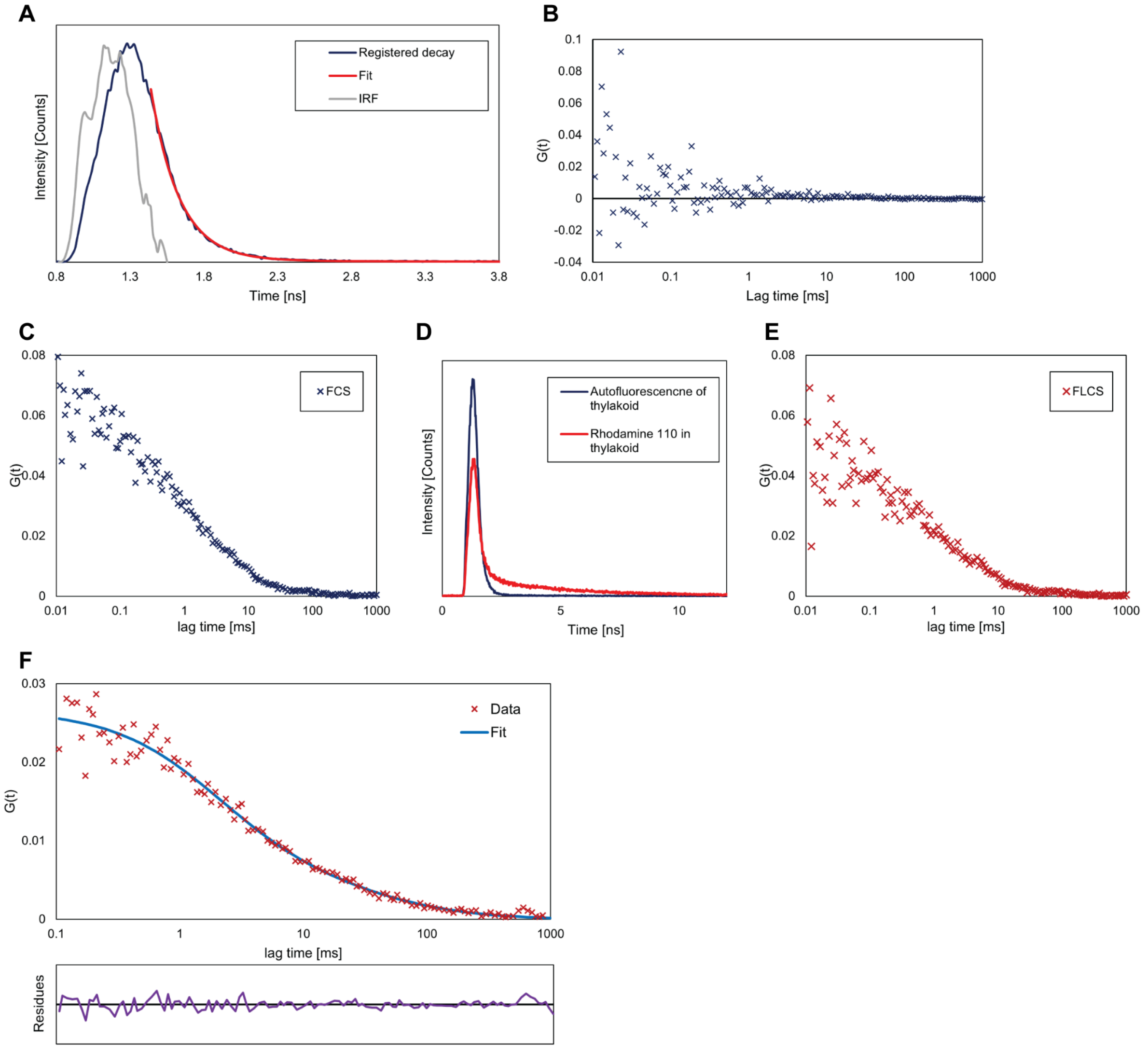


**Fig. S13. Fluorescence lifetime correlation spectroscopy (FLCS) removes thylakoid autofluorescence and isolates the diffusion signal of Rhodamine 110.** (A) Fluorescence lifetime decay of thylakoid autofluorescence under 485 nm pulsed laser excitation. The registered decay is shown together with the instrument response function (IRF) and the fitted decay (Fit). The autofluorescence lifetime obtained from fitting was 0.2 ns. (B) Autocorrelation curve of thylakoid autofluorescence intensity fluctuations. No correlation pattern characteristic of diffusion is visible, indicating that no mobile autofluorescent species are present in the thylakoid structure. (C) FCS curve obtained from the unprocessed fluorescence signal of Rhodamine 110 in thylakoids. The autocorrelation is affected by photons originating from thylakoid autofluorescence rather than the tracer. (D) Comparison of fluorescence lifetimes of Rhodamine 110 in thylakoids (red) and thylakoid autofluorescence alone (dark blue). FLCS filtering removes photons with lifetime characteristics of autofluorescence, retaining only those with longer lifetimes. (E) FLCS curve after autofluorescence filtering, containing only diffusion information for Rhodamine 110. (F) Representative FLCS curve of Rhodamine 110 diffusing in thylakoids (gyroid-containing sample) fitted with a two-component free diffusion model. Residuals are shown below.
